# LASSO: versatile and selective biomolecule pulldown with combinatorial DNA-crosslinked polymers

**DOI:** 10.1101/2025.07.24.666000

**Authors:** Sarah Speed, Krishna Gupta, Yu-Hsuan Peng, Elisha Krieg

**Affiliations:** Division of Polymer Biomaterials Science, Leibniz Institute of Polymer Research Dresden, Germany; Faculty of Chemistry and Food Chemistry, Technische Universität Dresden, Germany

**Keywords:** biomolecule pulldown, DNA nanotechnology, phase separation, programmable materials, RNA-seq

## Abstract

Current methods for sequence-selective biomolecule isolation suffer from high cost, off-target effects, and limited flexibility. Here, we introduce *LASSO* (cross**L**ink-**A**ssisted **S**equence-**S**elective is**O**lation), a versatile platform using programmable polymer phase separation to capture biomolecules under native conditions. LASSO relies on *combinatorial crosslinker libraries*—diverse mixtures of DNA strands that collectively trigger the formation of highly swollen polymer agglomerates with near-zero background binding. We demonstrate >80% pulldown efficiency for diverse targets, including DNA, SARS-CoV-2 RNA, and human thrombin. LASSO provides 8–20x higher binding capacity (4 nmol/mg polymer) than commercial microbeads. In RNA-seq workflows, LASSO depleted ribosomal RNA with 86% efficiency while yielding up to 7x fewer off-target outliers (p<0.001) versus state-of-the-art magnetic beads (*riboPOOLs*) and RNase H (*NEBNext*) kits. Thrombin was captured via switchable aptamers with 92% efficiency, and a gentle release mechanism allowed the subsequent isolation of 72% enzymatically active proteins from the polymer. LASSO’s cost-effectiveness ($0.96/sample vs. $46–$51 for commercial kits), long-term stability (7+ years), simple usage, and modularity position it to transform diagnostics, transcriptomics, and bionanotechnology workflows.

## Introduction

The separation and purification of biomolecules from complex mixtures is a fundamental process in biology, biotechnology, and diagnostics.^1^ Traditional pulldown methods use inexpensive solid-phase resins to separate complex biomolecule mixtures based on molecular size, charge, or other features. The resins bind broad categories of biomolecules (e.g., nucleic acids) while leaving others in solution.^2,3^ However, more specific methods are often needed to isolate biomolecules in a sequence-selective manner. This is typically accomplished through complementary sequence hybridization (for nucleic acids) or through antibodies (for proteins). Magnetic microbeads functionalized with affinity ligands have emerged as the dominant solution for sequence-selective target capture.^4–7^ After binding their target, the microbeads are readily pulled down with a magnetic field. Other popular approaches utilize enzymes to selectively degrade non-target nucleic acid sequences.^8,9^ The greatest drawbacks of currently available sequence-selective methods are their relatively high cost (e.g., due to the use of enzymes or other recombinant proteins), off-target effects (e.g., due to non-specific adsorption on microbeads), and time-consuming sample preparation.^10–12^

Over recent decades, several stimuli-responsive or “smart” polymers have been developed as a scalable option for bioseparation. Such polymers respond to small changes in environmental conditions, such as temperature, ionic strength, pH, or electric or magnetic fields.^13–15^ Specific ligands for target capture can be linked to the polymer backbone, forming an “affinity macroligand.”^16–20^ After the target is bound to this ligand, the polymer is precipitated, thereby pulling the target out of solution. We previously used this principle to develop a methanol-responsive polymer (MeRPy) that selectively binds and isolates single-and double-stranded DNA targets.^21,22^ After binding target sequences, the addition of methanol leads to the reversible precipitation of MeRPy and thus the rapid purification of captured DNA targets from complex mixtures.

Currently, three challenges limit the wider adoption of smart polymers in bioseparation: First, the stimulus that triggers precipitation of the polymer (e.g., temperature, pH, co-solvent) can cause target denaturation or the loss of its biological function. Second, polymer precipitation may be triggered inadvertently during target capture. For example, binding of nucleic acids to hybridization probes often requires thermal annealing, which would cause premature precipitation of thermoresponsive polymers. Third, the precipitating polymer can cause nonspecific co-precipitation of non-target biomolecules.^23,24^ For instance, while MeRPy pulldown is sequence-selective for DNA,^22^ it fails to achieve the same selectivity for RNA targets.

Here we report a versatile pulldown method termed LASSO (cross**L**ink-**A**ssisted **S**equence-**S**elective is**O**lation), which overcomes the common limitations of smart polymers in affinity precipitation. LASSO uses a bio-orthogonal crosslinking mechanism based on diverse combinatorial DNA libraries to trigger polymer agglomeration and phase separation under otherwise unchanged (native) conditions. This mechanism does not require organic solvents or other harsh environmental conditions. We show that LASSO can capture DNA, RNA, or proteins with high specificity and efficiency, as well as release selected targets back into solution. We demonstrate a practical application of LASSO in ribosomal RNA (rRNA) depletion for RNA sequencing (RNA-seq) library preparation, achieving efficient depletion with reduced off-target bias as compared to the two most widely used commercial rRNA depletion methods. We further demonstrate the pulldown and release of human thrombin protein under native conditions, preserving the protein’s structure and enzymatic function.

## Results

### Material concept

LASSO relies on three components: an acrylamide-based *polymer*,^21,22,25^ DNA oligonucleotide *catcher strands*, and DNA *crosslinkers* (Figure 1, Supplementary Figure 1). The polymer is functionalized with DNA *anchor strands*. These serve as binding sites to program the material’s phase separation and affinity characteristics with crosslinkers and catcher strands, respectively. We synthesized two variants of the polymer, **P**_**10**_ and **P**_**20**_, differing in their degree of functionalization with anchor strands (see Methods section, Supplementary Figures 2, 3). The catcher strands comprise a target-specific binding domain, an adapter domain complementary to the anchor strand, and an (optional) toehold domain for triggering target release through toehold-mediated strand displacement (TMSD) (Supplementary Figure 1a,c). The binding domain consists of either a linear DNA sequence to capture DNA or RNA via hybridization or an aptamer sequence to capture a protein.

**Figure 1.**
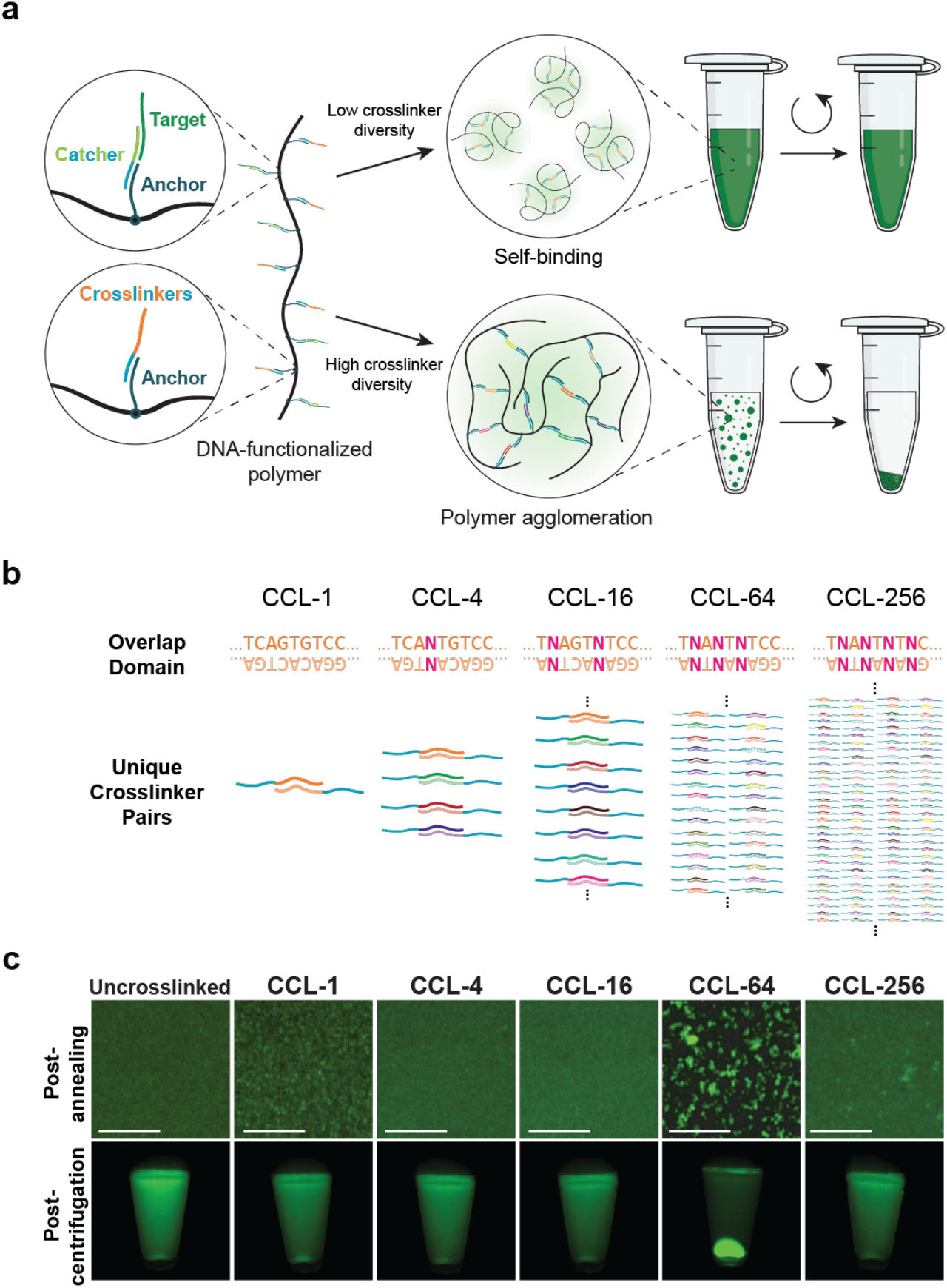
LASSO uses crosslinking-induced phase separation for selective biomolecule pulldown under mild conditions. **a)** Target-specific DNA catcher strands and DNA crosslinkers hybridize to DNA anchor strands conjugated to ultra-high molecular weight polymer chains. A low diversity of crosslinkers leads to ineffective intra-molecular bond formation, whereas a high diversity of crosslinkers promotes inter-molecular crosslinking and phase separation. Target-bound polymer agglomerates are centrifuged into a pellet, separating and encapsulating the target. **b)** Combinatorial crosslinker library (CCL) approach in which the overlap domain is diversified through the introduction of mixed “N” nucleotides. The complexity of the library increases with the number of N nucleotides in the sequence. **c)** Effect of CCL diversity on phase separation. **Top row:** Fluorescent microscope images of **P**_**10**_ without crosslinkers or crosslinked with different CCLs at 100x magnification and stained with a Cy5-labeled target DNA oligonucleotide. Scale bar: 15 µm. **Bottom row:** Pulldown of a Cy5-labeled target DNA oligonucleotide on **P**_**10**_ crosslinked with different CCLs.

We envisioned that the addition of DNA crosslinkers would trigger the formation of large polymer agglomerates, leading to phase separation and thereby providing a gentle pulldown mechanism (Figure 1a, Supplementary Figure 1b). Such a *crosslink-induced secondary effect affinity precipitation* is highly uncommon,^26,27^ likely because in dilute polymer solutions, intra-molecular binding is favored over crosslinking,^28,29^ making polymer agglomeration vastly inefficient (Figure 1a). However, we recently discovered that this problem can be solved by using *combinatorial crosslinker libraries* (CCL)—complex mixtures of DNA strands containing many distinct recognition sites.^29^ Sufficient crosslinker diversity suppresses ineffective intra-molecular bonds. We therefore followed our previous approach,^29^ in which the overlap domains of crosslinker pairs are diversified by introducing *mixed bases* (N) at specific positions, where N can be either of the four canonical nucleobases with roughly equal probability (Figure 1b). The number of unique crosslinker pairs in the CCL is 4^n^, where n is the number of N bases in the sequence.

We first tested how crosslinker diversity affected the phase separation of **P**_**10**_, using five different CCLs with n = 0 (“CCL-1”), n = 1 (“CCL-4”), n = 2 (“CCL-16”), n = 3 (“CCL-64”), and n = 4 (“CCL-256”). Notably, large polymer agglomerates only formed with CCL-64, but not with CCLs having substantially smaller or larger diversities (Figure 1c, top row). The existence of this surprising “Goldilocks zone” in sequence diversity is likely due to counteracting effects on binding topology versus binding kinetics and thermodynamics (Supplementary Note 1). The same trend was also observed in pulldown experiments using a benchtop centrifuge, where only CCL-64 crosslinked polymers pelleted efficiently to isolate a fluorescently labeled DNA target (Figure 1c, bottom row; Supplementary Procedure 1).

LASSO’s CCL-64-crosslinked polymers form a sparse molecular network that is swollen by 99.7% water (w/v), with an estimated mesh size (ξ) of 32 nm, thus allowing efficient permeation with both small and large biomacromolecular targets (see Methods section). The absence of a sharp solid-liquid interface, and the low polarizability contrast with the medium disfavor non-specific surface adsorption.^30^ These properties were expected to provide substantially increased binding capacity and selectivity as compared to solid resins or microbeads.

We optimized LASSO by testing the pulldown of DNA using different concentrations of **P**_**10**_ and CCL-64 (Supplementary Figures 4 and 5, Supplementary Note 2). The most efficient target capture was observed when using a polymer concentration of 0.05% (w/v), while 80% of available anchor strands were bound to CCL-64 and 10% of anchors were bound to catcher strands. Under these conditions, LASSO can bind 2 nmol target per milligram **P**_**10**_ and 4 nmol target per milligram **P**_**20**_.

### LASSO efficiently and specifically captures DNA and RNA targets

To demonstrate the efficiency and specificity of the LASSO method for biomolecule capture, we used CCL-64-crosslinked **P**_**10**_ to capture single-stranded DNA and RNA (Supplementary Procedure 2). A 22-nucleotide (nt) DNA target was captured with an average efficiency of 81.2 ± 0.1%. Control samples lacking either the catcher strands or the crosslinkers showed no pulldown (Figure 2a,b), highlighting the high capture specificity. Next, we tested the ability of LASSO to capture SARS-CoV-2 N-gene RNA (1442 nt) to demonstrate potential application in sequestration of viral RNA for diagnostic assays. Each supernatant was subjected to RT-qPCR to determine the amount of remaining N-gene RNA (Figure 2c, Supplementary Figure 6). On average, LASSO captured N-gene RNA with an efficiency of 92.3 ± 1.5% (Figure 2c). In the absence of catcher strands, a pellet formed, but no significant target capture was observed, confirming a high binding specificity for the long RNA target.

**Figure 2.**
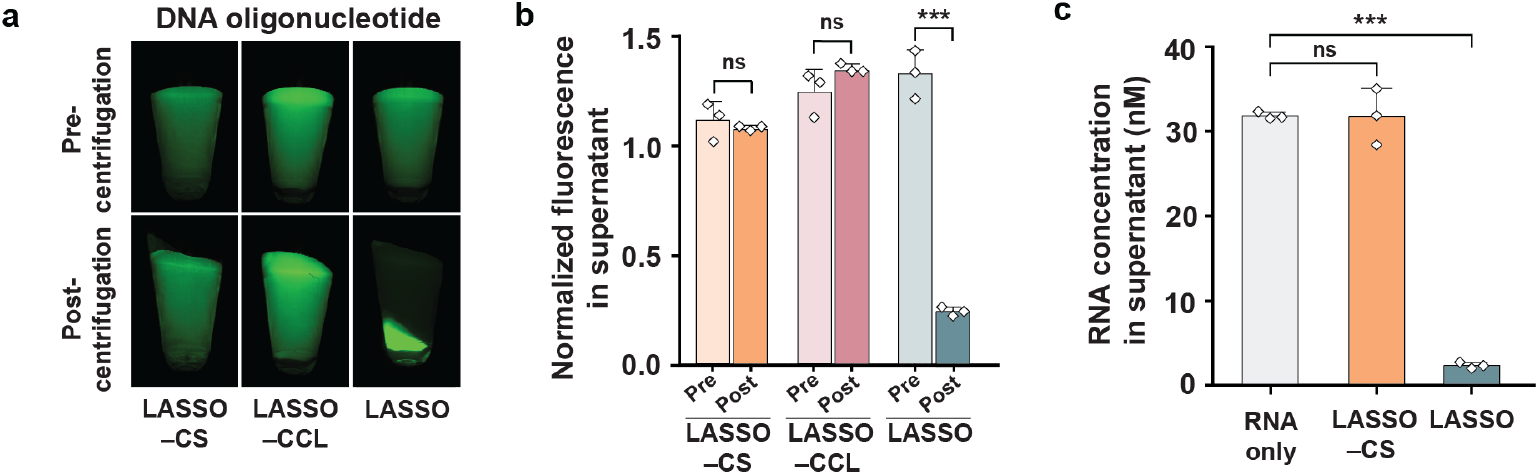
Programmable nucleic acid capture with LASSO is efficient and highly specific. **a)** Capture and pulldown of a fluorescent DNA oligonucleotide is only observed in the presence of both catcher strands (CS) and crosslinkers (CCL). **b)** Quantification of the DNA target remaining in the supernatant after pulldown, based on fluorescence intensity. **c)** Concentration of SARS-CoV-2 N-gene RNA in the supernatant of the indicated samples as determined by RT-qPCR (Supplementary Figure 6). Data for panels b and c are shown as mean ± s.d. (*n* = 3 independent experiments). Statistical analysis was performed using an unpaired two-tailed t-test; ns, non-significant (p > 0.05); \*\*\**p* < 0.001.

### LASSO enhances RNA-seq expression profiles

We next explored whether LASSO can be used to enhance libraries for RNA-seq. Most library preparation methods involve the removal of highly abundant and non-informative transcripts such as rRNA to increase the sensitivity for functionally more relevant transcripts. The depletion of abundant species should ideally not affect non-target transcripts or otherwise bias the transcriptomic profile.

We benchmarked LASSO against two of the most widely used commercial rRNA depletion kits, *riboPOOLs* and *NEBNext*. These two kits represent the two prevailing approaches for target depletion: riboPOOLs is based on magnetic bead capture, whilst NEBNext relies on enzymatic target degradation with RNase H.^8,9,31–34^ We surmised that LASSO may offer superior specificity, since beads are prone to nonspecific interfacial adsorption,^10,12^ and the enzyme-based assays can inadvertently degrade transiently bound non-target transcripts.^11^ Both cases can lead to undesired loss of non-target transcripts and systematically biased transcriptomic data.

We first generated an rRNA-specific *catcher strand library* (CSL) containing 195 distinct catcher strands that tile across the sequences of all six human rRNAs, namely, cytoplasmic rRNAs (28S, 18S, 5.8S, and 5S) and mitochondrial rRNAs (12S and 16S). Each catcher strand contains a 45–50-nt target-binding domain^35^ and a 20-nt adapter domain (Supplementary Data 1, strand IDs 22–216). The **P**_**20**_ backbone was used instead of **P**_**10**_ to ensure that many different catcher strands could be captured at sufficiently high individual concentrations. We then compared the performance of LASSO (Supplementary Procedure 3), riboPOOLs, and NEBNext for the depletion of rRNA from human total RNA. A control sample treated with LASSO in the absence of CSL (LASSO –CSL) was also included for reference.

All treated samples as well as the original total RNA were sequenced to compare transcriptomic data before and after rRNA depletion. The proportion of total RNA reads that mapped to rRNA was 76.0 ± 1.4% before pulldown and 75.4 ± 0.2% for LASSO –CSL. When programmed with the CSL, LASSO reduced the number of rRNA reads to 10.7 ± 5.7%, corresponding to a pulldown efficiency of ∼86%. riboPOOLs and NEBNext demonstrated higher depletion efficiencies, reducing rRNA reads to 2.3 ± 0.3% and 0.1 ± 0.1%, respectively (Figure 3a, Supplementary Data 2). While the newly purchased NEBNext kit showed the highest pulldown efficiency, an expired kit yielded drastically reduced rRNA depletion, with 66% of rRNA remaining in the sample (Figure 3a). This outlier was excluded from further analysis, but the strong decrease in depletion efficiency highlights the susceptibility of enzyme-based assays to degradation and variations in performance.

**Figure 3.**
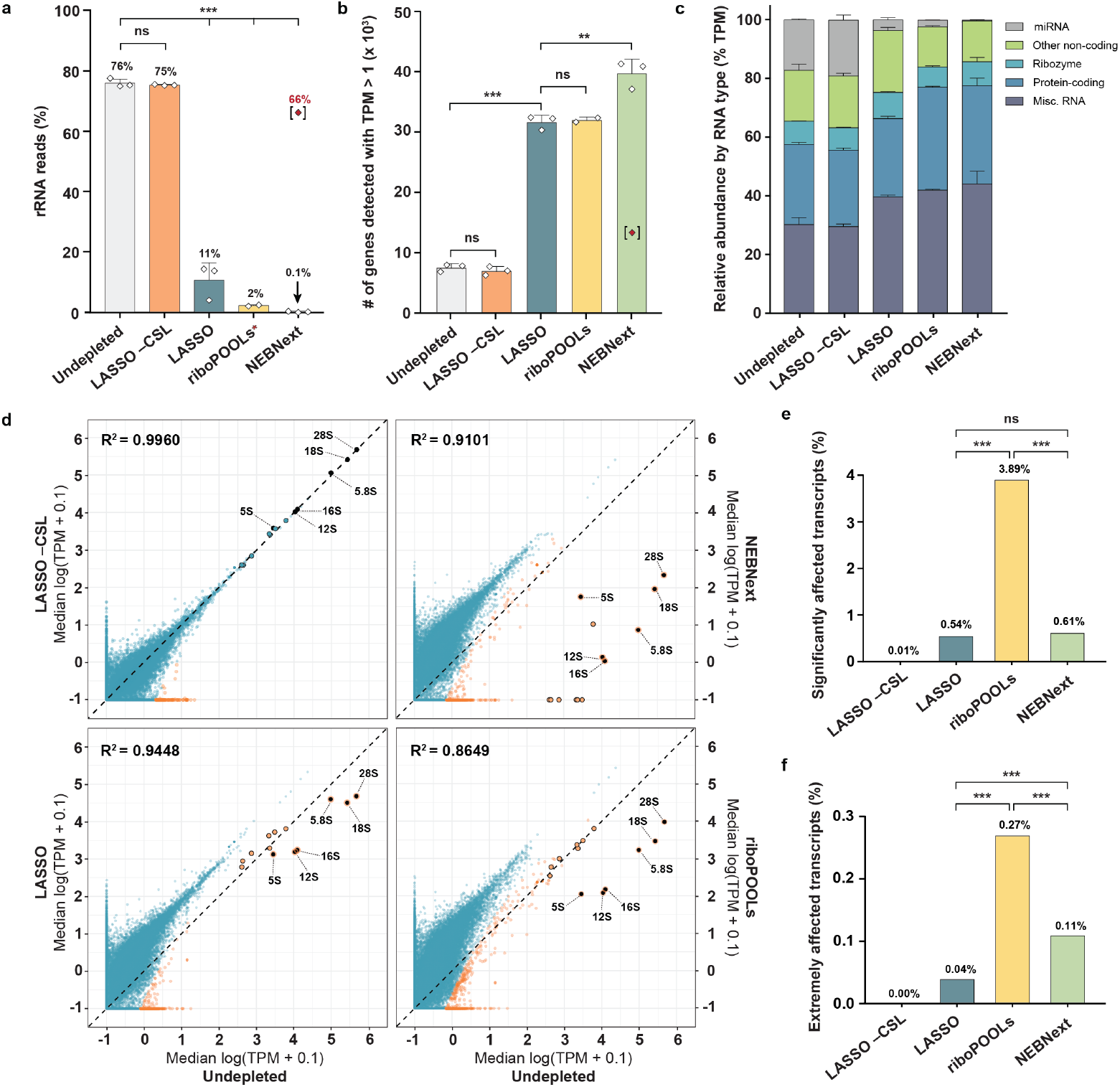
LASSO introduces fewer biases than commercial rRNA depletion kits for RNA-seq, while providing similar increases in sequencing depth. **a)** Percentage of RNA-seq reads mapped to rRNA for each condition. The red asterisk for the riboPOOLs data indicates the failure of one replicate to sequence properly due to the low final concentration. The bracketed red datapoint for the NEBNext condition corresponds to a sample prepared with an expired kit, which was excluded from data analysis. **b)** Total number of transcripts detected with >1 TPM for each condition. Statistical analysis for panels a and b was performed via an unpaired two-tailed t-test. **c)** All non-rRNA reads were mapped to their respective Ensembl biotype annotation, normalized to transcripts per million (TPM), and grouped by type. **d)** Correlation analysis of expression profiles between rRNA-depleted conditions and the undepleted condition. The six targeted rRNA transcripts are labeled. Orange data points represent outliers whose residuals lie >3 s.d. away from the linear trendline (see Methods section). Data points with black outlines indicate transcripts with high sequence similarity (>90%) to rRNA. **e)** Percentage of significantly affected and **f)** extremely affected transcripts for each method as shown in the volcano plots in Supplementary Figure 11. Transcripts with padj < 0.05 and absolute log_2_ fold change > 1 or padj < 0.001 and absolute log_2_ fold change > 3 are considered significantly or extremely affected, respectively. Statistical analysis was performed using a two-sided Fisher’s exact test. All data in panels a, b, and c are shown as the mean ± s.d.; data in panel d is shown as the median (n = 2 independent experiments for the riboPOOLs condition; n = 3 independent experiments for all other conditions). ns, non-significant (p > 0.05); \*\**p* < 0.01; \*\*\**p* < 0.001.

For all three methods, rRNA depletion significantly increased the number of reads exceeding the threshold of one transcripts per million (TPM) (Figure 3b, Supplementary Figure 7). LASSO depletion provided a 4.3 ± 0.5-fold increase in TPM values >1 for non-rRNA transcripts, demonstrating a substantial enhancement of the RNA-seq library. The increase in TPM values >1 was statistically indistinguishable from riboPOOLs (4.4 ± 0.4-fold), but lower than for samples depleted by the newly purchased NEBNext kit (5.3 ± 0.1-fold).

To compare undesirable off-target depletion between the three methods, we first mapped all non-rRNA reads in each library to the Ensembl RNA biotypes (Figure 3c, Supplementary Figure 8, Supplementary Data 3). Among the three tested methods, depletion with LASSO produced the least biased biotype distribution, retaining a significantly higher percentage of micro RNAs (miRNA) and small nuclear RNAs (snRNA; included in the “other non-coding” biotype), when compared to NEBNext (p = 0.0018 for miRNA; p = 0.0004 for snRNA) and riboPOOLs (p = 0.0034 for snRNA) (Figure 3c).

We also correlated the expression levels of individual transcripts in the depleted samples with those of the original undepleted samples (Figure 3d, Supplementary Figure 9). Pearson correlation coefficients (R) were highest for LASSO (R = 0.959–0.977), followed by NEBNext (R = 0.934–0.955) and riboPOOLs (R = 0.919–0.934) (Supplementary Figure 9). To highlight systematic biases arising from non-target depletion, we marked transcript outliers in the expression plots (Figure 3d, orange data points, Supplementary Figure 10; see Methods section). The significance of the outliers was further quantified by differential analysis of transcript counts (Figure 3e,f, Supplementary Figure 11). Despite their widespread use, substantial depletion of several non-rRNA transcripts were observed in both the NEBNext and riboPOOLs samples. riboPOOLs produced the largest number of significantly affected and extremely affected outliers (3.89% and 0.27% of transcripts, respectively). NEBNext produced fewer outliers than riboPOOLs (0.61% significant and 0.11% extreme of transcripts). However, several abundant non-target transcripts were depleted by more than 3 orders of magnitude or even entirely eliminated from the library (Figure 3d, Supplementary Figure 11). LASSO produced the lowest number of outliers out of all three methods (0.54% significant and 0.04% extreme outliers) (Figure 3e,f, Supplementary Figure 11).

Among the highly expressed transcripts (with original TPM >100), LASSO produced only seven outliers, compared with 49 outliers for riboPOOLs and 35 outliers for NEBNext. All seven outliers in the LASSO samples were also affected in the commercial methods, and in some cases completely removed in the NEBNext samples. Further analysis revealed that these seven common outliers are rRNA-derived miRNAs and long non-coding RNAs (lncRNA) that share >90% similarity with the targeted rRNA sequences (Figure 3d, data points with black outlines, Supplementary Data 4). Among these outliers are MIR3648-1/-2, MIR663A/B, and MIR663AHG, which have been previously associated with oncogenesis in gastric, colon, and kidney cancers.^36–38^ Other highly expressed transcripts such as RNU2-1 and the U2 family of snRNAs were undesirably depleted in riboPOOLs and NEBNext, but fully retained in LASSO. These RNAs are essential components of the spliceosome and are overexpressed in lung, pancreatic, and colorectal cancers, making them potential candidates as diagnostic biomarkers in serum.^39,40^

Notably, LASSO –CSL showed no significant distortions in biotype distribution and virtually no individual transcript outliers (Figure 3c,f; Supplementary Figure 11). Therefore, all instances of non-specific pulldown in LASSO can be attributed to spurious interactions of the transcripts with the CSL. This data demonstrates that the phase separating polymer itself does not interfere with any components of the transcriptome, thus providing an exceptional degree of near-zero background capture specificity.

### Switchable aptamers enable protein capture and release

To further demonstrate the broad utility of LASSO, we programmed **P**_**10**_ to capture and release human thrombin with a switchable aptamer CSL (Figure 4a, Supplementary Procedure 4). To this end, we created switchable versions of two previously reported thrombin aptamers, TBA and HD22,^41,42^ by including a toehold sequence downstream of the aptamer domain of the catcher strand (Supplementary Figure 1c). The addition of a suitable release strand was expected to trigger a TMSD reaction,^43,44^ thereby switching the aptamer from a folded accessible state to a fully double-stranded inaccessible state. This switching mechanism would release the captured native protein back into solution (Figure 4a).

**Figure 4.**
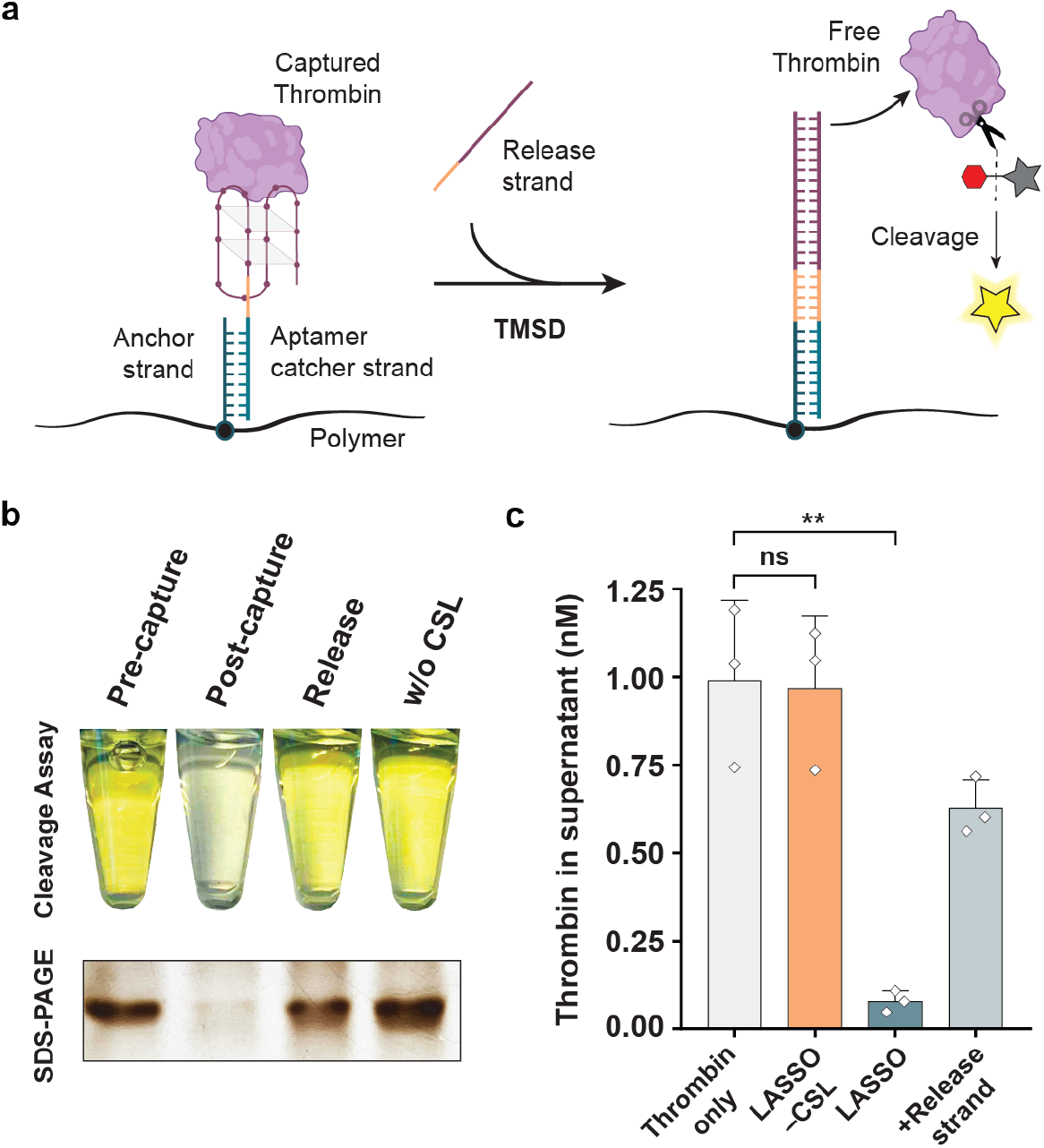
LASSO gently captures and releases thrombin through switchable aptamer binding. **a)** Scheme depicting the release of captured thrombin by the addition of a release strand that switches the aptamer strand from an accessible folded state to an inaccessible double-stranded state through TMSD. **b)** Cleavage assay in which S-2238 substrate changes from colorless to yellow through selective cleavage by thrombin in the supernatant. The SDS-PAGE gel below shows the relative amounts of thrombin present in the corresponding samples above. The original tube and gel images are shown in Supplementary Figure 12. **c)** Concentration of thrombin in the supernatant of the indicated samples as determined by absorbance measurements of the cleaved substrate. Data are shown as mean ± s.d. (*n* = 3 independent experiments). Statistical analysis was performed using an unpaired two-tailed t-test; ns, non-significant (p > 0.05); \*\**p* < 0.01.

To test both the capture efficiency and the activity of thrombin after release, we performed a cleavage assay with the thrombin-specific chromogenic peptide substrate S-2238. This experiment allowed for quantitative colorimetric analysis of thrombin activity in the supernatant before and after capture, and after its release back into the solution (Figure 4b,c; Supplementary Figure 12). We additionally confirmed the presence of thrombin by SDS-PAGE (Figure 4b, Supplementary Figure 12). We observed a significant decrease in thrombin concentration and activity in the supernatant after capture on LASSO, corresponding to a 91.9 ± 3.1% capture efficiency. After the addition of the release strand, 71.7 ± 19.3% of the captured thrombin was released back into the supernatant. No significant capture was observed in the absence of the aptamer CSL (Figure 4c). These results demonstrate that LASSO enables gentle and selective capture and release of active proteins in their native form.

## Discussion

LASSO is a programmable and selective method for biomolecule pulldown. Its key feature is a novel phase separation mechanism that relies on combinatorial crosslinker libraries (CCL): mixtures of many different DNA strands that generate highly permeable polymer agglomerates that can scavenge target molecules and sediment during centrifugation. This approach complements recent advancements in phase-separation engineering based on nucleic acids.^45–47^ Importantly, the entire capture and separation process occurs in free solution, without the involvement of solid-liquid interfaces. This reduces the likelihood of nonspecific surface adsorption, which is a persistent challenge for methods based on microbeads and other solid substrates.^10,12^ The ability to capture targets in a homogenous phase also enables very high binding capacities of up to 4 nmol per milligram polymer, while bead-based assays typically bind 0.2–0.5 nmol per milligram substrate.^48^

LASSO addresses historical shortcomings of smart polymers and circumvents many drawbacks of earlier approaches in affinity precipitation.^19,26,49^ Unlike other DNA-functionalized polymer systems,^26,27,49^ such as the popular thermo-responsive poly(N-isopropylacrylamide),^18^ LASSO pulldown does not require heating-induced precipitation, addition of chemicals, or other harsh environmental changes to capture targets. Therefore, the polymer can be programmed to isolate diverse types of biomolecules sequence-selectively and under native conditions, which is ideal for sensitive targets. We achieve consistently high capture efficiencies (81–92%) for DNA, RNA, and protein in a one-pot reaction that requires only basic laboratory equipment (Figure 5). The ability to release target proteins was demonstrated by using a pair of switchable aptamers, but the underlying displacement mechanism can be easily applied to any other target bound to DNA-based catcher strands.^22^ The versatility of this method represents a significant advancement over previously reported MeRPy pulldown,^21,22,50^ which is applicable for DNA capture but degrades proteins and non-specifically binds RNA (Supplementary Figure 13).

**Figure 5.**
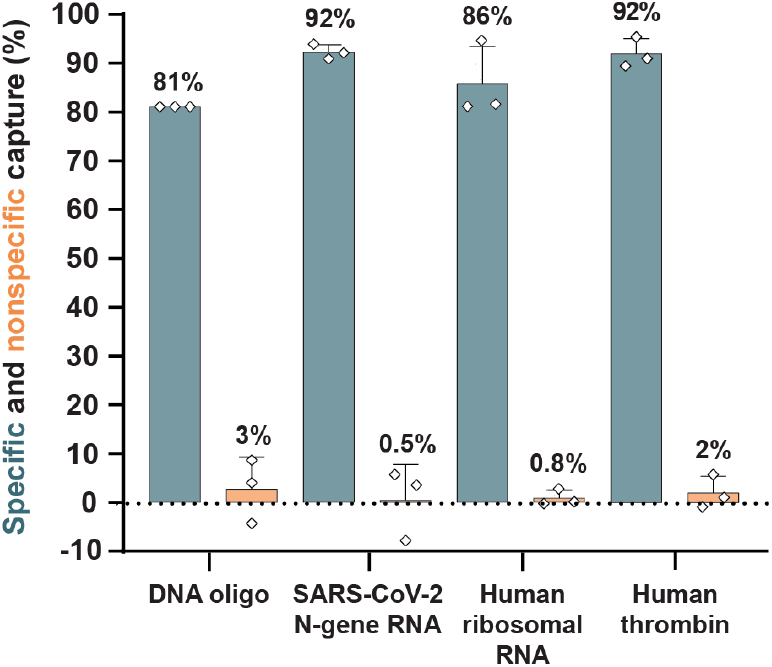
Summary of biomolecule capture on LASSO. LASSO enables highly specific and efficient capture of DNA, RNA, and protein in a programmable and target-specific manner. Capture efficiencies were quantified by fluorescence measurements (DNA oligo), RT-qPCR (SARS-CoV-2 N-gene RNA), RNA-seq (human ribosomal RNA), and absorbance measurements (human thrombin). Nonspecific capture was measured from samples subjected to LASSO pulldown without CSL under otherwise identical conditions. The data are shown as mean ± s.d. (*n* = 3 independent experiments).

Constructing and purifying sequencing libraries is one of the most immediate applications for LASSO. We depleted rRNA from RNA-seq libraries, substantially enhancing transcriptomic profiles while demonstrating minimal off-target bias. In particular, non-coding RNAs, which are of interest for the study of disease biomarkers and gene regulation,^51,52^ were more effectively retained in the LASSO-depleted samples as compared to the commercial kits. We suspect that the biases introduced in riboPOOLs are predominantly caused by nonspecific surface adsorption, which would explain the numerous off-target outliers that do not have substantial sequence similarity to rRNA. In contrast, biases created by the NEBNext kit are attributed to its *irreversible* depletion mechanism: if a non-target RNA molecule binds to a probe, even transiently, it can get digested by RNase H, and is thereby irreversibly lost from the library. This explanation is consistent with the extremely strong off-target depletion of certain miRNAs and lncRNAs that have high sequence similarity with rRNA. The distinctly different mechanism of LASSO circumvents both sources of bias: off-target effects are limited to sequences similar to rRNA and their depletion is weaker than the rRNA pulldown itself.

By requiring only synthetic polymer components, LASSO also provides higher stability and much lower cost when compared to conventional methods (Supplementary Tables 1 and 2). Depleting rRNA from 1 µg of total RNA with LASSO costs approximately $0.96 per sample, while the two commercial kits cost $46 to $51 per sample. Storage and transportation of LASSO components do not require a cold chain, making it suitable for applications in resource-limited settings. We further found that the DNA-functionalized polyacrylamide backbone can be stored in the freezer for at least 7 years, without measurable reduction in binding capacity (Supplementary Figure 14).

Overall, LASSO enables reliable capture of diverse biomolecular targets while providing exceptional pulldown specificity and the ability to recover targets under gentle conditions. Depletion of rRNA from RNA-seq libraries with LASSO increases sequencing depth while introducing significantly lower biases in the transcriptomic data than state-of-the-art commercial kits. Our future studies will explore the use of this method for streamlining point-of-care diagnostic tests.^53,54^ In particular, LASSO could be used to enrich low abundance biomarkers from clinical samples in a simple one-step procedure. We anticipate that LASSO will improve preparative workflows across diverse applications, including diagnostics, bionanotechnology, next-generation sequencing, and transcriptomics.

## Methods

### Materials

Solvents and reagents were purchased from commercial sources and used as received, unless otherwise specified. Water was obtained from a Milli-Q system from Merck Millipore. Molecular biology grade acrylamide (catalog number A9099) and 19:1 acrylamide/bis-acrylamide (catalog number A2917), sodium acrylate (catalog number 408220), and ammonium persulfate (APS; catalog number A3678) were purchased from Sigma-Aldrich. Methanol (ACS reagent grade; catalog number 423955000), ultrapure N,N,N′,N′- tetramethylethylenediamine (TEMED; catalog number 15524010), and SYBR™ Gold Nucleic Acid Gel Stain (catalog number S11494) were purchased from Thermo Fisher Scientific. Desalted oligonucleotides were purchased from Integrated DNA Technologies (IDT). Nitrogen gas (>99.999%) was used under inert conditions and supplied by an in-house gas generator. To ensure an inert condition, nitrogen gas was purified through a Model 1000 oxygen trap from Sigma-Aldrich (catalog number Z290246). Reagents with unreacted acrylamide groups were stored at 4 °C or −20 °C, protected from unnecessary exposure to light. The NEBNext^®^ rRNA Depletion Kit v2 with Beads (catalog number E7405S), Monarch^®^ Spin RNA Cleanup Kit (catalog number T2050S), and DNase I (catalog number M0303S), were purchased from New England Biolabs (NEB). The riboPOOLs rRNA depletion kit with cleanUP module was purchased from siTOOLs Biotech (catalog number dp-K012-53). Human HeLa cell total RNA was purchased from Takara Bio (catalog number 636543) and stored at −80 °C until use. Purified thrombin from human plasma (catalog number T6884) and bovine serum albumin (BSA, catalog number A2934) were purchased from Sigma-Aldrich. Chromogenix S-2238™ thrombin substrate was purchased from Diapharma (catalog number S820324). DNA ladders (catalog number SM1211) and protein ladders (catalog number LC5615) were purchased from Thermo Fisher Scientific.

### Polymer synthesis

A detailed protocol is available in ref.^22^. In brief, acrylamide (50 mg ml^−1^), sodium acrylate (0.5 mg ml^−1^), and acrylamide-labeled anchor strand DNA (Supplementary Data 1, strand ID 1) were co-polymerized at a molar ratio of 10,000:100:x (x = 10 or 20 for **P**_**10**_ and **P**_**20**_, respectively) in 1x TBE buffer (100 mM Tris, 90 mM boric acid, 1 mM EDTA, pH 8.3) to create DNA-grafted poly(acrylamide-coacrylic acid). Synthesis of **P**_**10**_ was initiated by addition of 0.025 wt% TEMED and 0.025 wt% APS. Synthesis of **P**_**20**_ was initiated by addition of 0.05 wt% TEMED and 0.05 wt% APS. To achieve high molecular weight and a narrow size distribution, it was necessary to carry out the reaction in high-purity nitrogen gas, which was passed through an oxygen trap on-site. The reaction was allowed to proceed overnight, resulting in a highly viscous polymer solution indicating the formation of long polymer chains. NMR spectroscopy was used to verify high conversion of the monomer (Supplementary Figure 2). The solution was diluted in 9 volumes of 1x TE buffer (10 mM Tris, 1 mM EDTA, pH 8.0) and subsequently purified via methanol precipitation. The pellet was resuspended in milliQ water at 2.5% (w/v) and stored in aliquots at −20 °C. The binding capacities of **P**_**10**_ and **P**_**20**_ were found to be 15 nmol single-stranded DNA (ssDNA) per milligram polymer and 34 nmol ssDNA per milligram polymer, respectively (Supplementary Figure 3).

### Polymer mesh size calculation

To estimate the mesh size (ξ) of the polymer network, we applied the relationship ξ ≈ ν_e_^−1/3^, where ν_e_ is the effective crosslinker concentration in units of molecules per cubic meter, which is derived from the affine network model of rubber elasticity.^55^ Based on the measured volume of the pellet after centrifugation, the crosslinker concentration in the sedimented polymers agglomerates was estimated to be 50 μM (ν_e_ = 3.01 × 10^22^ molecules/m^3^). Assuming a uniform, isotropic 3D network, the mesh size was calculated to be ξ ≈ (3.01 × 10^22^)^−1/3^ ≈ 32 nm. This value represents the average distance between effective crosslinks within the molecular network for **P**_**10**_.

### LASSO capture

Detailed step-by step protocols for biomolecule capture with LASSO are described in Supplementary Procedures 1–4. In brief, samples were prepared at a final concentration of 0.05% (w/v) **P**_**10**_ or **P**_**20**_, mixed with up to 0.1 molar equivalents of CSL to the anchor strand, 0.8 molar equivalents of CCL-64 to the anchor strand, and 0.2–0.3 molar equivalents of the target to the CSL in a buffer condition of 150 mM NaCl and 1x TE buffer (10 mM Tris, 1 mM EDTA, pH 8.0). The samples were annealed on a C1000 Touch™ Thermal Cycler (Bio-Rad) using the following steps: (1) heating at 95 °C for 3 min, (2) instant cooling from 95 °C to 80 °C, (3) holding at 80 °C for 2 min, (4) first cooling ramp from 80 °C to 65 °C at −1.5 °C min^−1^, (5) second cooling ramp from 65 °C to 37 °C at −2.8 °C min^−1^, (6) holding at 4 °C or at 20 °C until use. The first cooling ramp facilitates binding of the adaptor domains of the crosslinkers to the anchor strands, while the second ramp allows proper binding of the overlap domains to their complementary partners. For protein capture, the protein was added after annealing and 0.1% (w/v) BSA was added to prevent non-specific protein interactions. The protein-containing mixture was then incubated with light shaking for 30 minutes at room temperature. Samples were centrifuged for 30 minutes at 17,000 xg at 4 °C or at room temperature to pellet the target-bound polymer.

### SARS-CoV-2 N-gene RNA synthesis

A T7 promoter was added to the SARS-CoV-2 N-gene sequence through PCR amplification from a 2019-nCoV_N_Positive Control plasmid (IDT, catalog number 10006625) using the Q5^®^ Hot Start High-Fidelity 2X Master Mix (NEB, catalog number M0494S) and SAR-CoV-2 N-gene T7 primers (Supplementary Data 1, strand IDs 15 and 16).^54^ The PCR product was purified with a DNA Clean & Concentrator spin column kit (Zymo Research, catalog number D4033) and the concentration was quantified by UV/Vis absorbance data using an Implen NanoPhotometer^®^ P360. SARS-CoV-2 N-gene RNA was synthesized from the purified DNA product using a HiScribe^®^ T7 High Yield RNA Synthesis Kit (NEB, catalog number E2040S). Remaining DNA was digested by addition of 20 U ml^−1^ DNase I (NEB, catalog number M0303S). EDTA was added to a final concentration of 5 mM, and DNase I was subsequently inactivated by incubation at 75 °C for 10 minutes. The final RNA product was purified using a Monarch^®^ Spin RNA Cleanup Kit (NEB, catalog number T2050S). RNA quality and concentration were measured on a Bioanalyzer 2100 capillary electrophoresis system (Agilent) using the Agilent RNA 6000 Nano Kit (catalog number 5067-1511).

### DNA oligonucleotide capture analysis

Following capture of fluorescent DNA oligonucleotide on LASSO (Supplementary Procedure 2), fluorescence images of Eppendorf tubes (in the Cy5 channel; excitation at 635 nm) were recorded before and after centrifugation on a Typhoon FLA 9500 scanner (GE Healthcare Life Sciences) at a 50 µm pixel size with the accompanying software (v1.0). Supernatant samples before and after centrifugation were quantified in a Tecan Spark^®^ microplate reader (Tecan) by Cy5 dye fluorescence (excitation at 649 nm; emission at 697 nm).

### SARS-CoV-2 N-gene RNA capture analysis

Following capture of SARS-CoV-2 N-gene RNA on LASSO (Supplementary Procedure 2), supernatant samples were diluted 1:5000 in nuclease-free H_2_O supplemented with 1 U μl^−1^ RNase inhibitor (NEB, catalog number M0307S). RNA remaining in the supernatant was amplified via RT-qPCR using the Luna^®^ Universal One-Step RT-qPCR kit (NEB, catalog number E3005S) and SARS-CoV-2 N-gene RNA primers (Supplementary Data 1, strand IDs 20 and 21). A standard curve was prepared from purified SARS-CoV-2 N-gene RNA. The samples were thermocycled on a CFX96 C1000 Touch™ Real Time Thermal Cycler (Bio-Rad). Data were analyzed on Bio-Rad CFX Maestro software (v4.1.2433.1219).

### RNA sequencing (RNA-seq)

#### RNA-seq library preparation and sequencing

Human HeLa cell total RNA (Takara Bio, catalog number 636543) was subjected to rRNA depletion with i) LASSO (Supplementary Procedure 3), ii) the riboPOOLs rRNA depletion kit with cleanUP module (siTOOLs Biotech, catalog number dp-K012-53), and iii) the fresh or expired NEBNext rRNA Depletion Kit v2 (NEB, catalog number E7400S). The expired NEBNext rRNA Depletion Kit v2 was 14 months past the stated expiration date. The rRNA-specific CSL was adapted from ref.^35^ for use with LASSO. The procedures for each of the methods were carried out as independent triplicates, but one of the riboPOOLs replicates failed to pass quality control prior to sequencing, due to low final RNA concentration. NaCl was removed from the LASSO-depleted sample using a Monarch^®^ Spin RNA Cleanup Kit (NEB, catalog number T2050S), then further treated with 57 U ml^−1^ DNase I (NEB, catalog number M0303S) to digest any remaining oligonucleotides. EDTA was added to a final concentration of 5 mM and the final RNA library was purified using a Monarch^®^ Spin RNA Cleanup Kit. The commercially-depleted samples were purified according to manufacturer’s instructions using the included SPRI beads. The concentration of purified RNA samples was quantified with the QuantiFluor^®^ RNA System (Promega, catalog number E3310) in a Tecan Spark^®^ microplate reader (Tecan). Non-treated total RNA was submitted for sequencing as an undepleted control. Final libraries were subjected to 150-bp paired-end sequencing on the Illumina NovaSeq System (Eurofins Genomics). Each of the sequencing data contained ≥ 4.7 million reads (Supplementary Data 2).

#### Data analysis

Quality control, read trimming, and pseudo-alignment with Kallisto^56^ were performed within the nf-core/rnaseq pipeline (v3.14.0)^57^ to produce TPM values. The source *Homo sapiens* genome GRCh38 release 112 was acquired from Ensembl.^58^ To correct for incomplete rRNA annotation in the source genome, a gene annotation file acquired from the UCSC Genome Browser^59^ for GRCh38 was filtered for rRNA repeat annotations (see https://github.com/zxl124/rRNA_gtfs) and concatenated to the source file. Percentage of raw rRNA reads was quantified using Ribodetector (v0.3.1).^60^ RNA biotypes were manually annotated to the Kallisto TPM output using the biotype entries from Ensembl for GRCh38 to determine relative abundances per RNA type. The “other non-coding” category includes artifact, lncRNA, scaRNA, scRNA, snoRNA, snRNA, sRNA, TEC (“to be experimentally confirmed”), tRNA, vault RNA, and gene and pseudogene biotypes. The TPM expression plots and the heatmap of the Pearson correlation matrix were generated in R (v4.4.1), using ggplot2 and pheatmap packages. Outliers in the TPM expression plots were calculated in R (v4.4.1) as follows. The standard deviations of the LASSO –CSL log(TPM) values as a function of the undepleted log(TPM) values were determined by assuming expected values of x = y. These standard deviations were assumed to reflect the expected spread in RNA-seq data due to sample handling as well as the limitations of sequencing depth, especially at low read numbers. These standard deviation values were plotted against the undepleted log(TPM) values and an exponential model was fit to the data to obtain continuous values of expected standard deviation for all undepleted log(TPM) values (Supplementary Figure 10). For each TPM expression plot, a linear model was fit to the data and the expected standard deviations from the linear fit were calculated with the exponential model. Residuals greater than 3 standard deviations were marked as outliers. Differential transcript detection was assessed using DESeq2^61^ and plotted in volcano plots showing statistically significant (padj < 0.05 and absolute log_2_ fold change > 1) and extremely statistically significant (padj < 0.001 and absolute log_2_ fold change > 3) transcripts.

### Thrombin cleavage assay

Following the capture and release of thrombin on LASSO (Supplementary Procedure 4), the concentration of thrombin in the supernatant was determined by absorbance change produced from the cleavage of the chromogenic thrombin substrate Chromogenix S-2238™ (Diapharma, catalog number S820324) as follows. A standard curve was prepared from purified thrombin (Sigma-Aldrich, catalog number T6884). To each experimental and standard curve sample, 125 μM of Chromogenix S-2238™ was added, and absorbance change at 405 nm was monitored for 20 minutes at 37 °C in a Tecan Spark^®^ microplate reader (Tecan). The kinetic curve for each sample was reduced to the mean slope of OD min^-1^ and plotted on the standard curve to determine the concentration of thrombin in each sample.

### Sodium dodecyl sulfate-polyacrylamide gel electrophoresis (SDS-PAGE)

SDS-PAGE gels were prepared using a 30% (w/w) acrylamide/bis-acrylamide stock (37.5:1; Bio-Rad, catalog number 1610158). The supernatant samples following thrombin capture were denatured at 95 °C for 5 minutes in 1x SDS loading dye (62.5 mM Tris-HCl, 10% (v/v) glycerol, 2% (w/v) SDS, 0.01% (w/v) bromophenol blue, 5% (v/v) DTT, pH 6.8) and loaded into a 12.5% SDS-PAGE gel. The gel was run in 0.5x TBE buffer (50 mM Tris, 45 mM boric acid, 0.5 mM EDTA, pH 8.3) at 100V on a XCell SureLock Mini-Cell Electrophoresis System using a Consort™ EV265 Electrophoresis power supply (Thermo Fisher Scientific). The gel was stained with the Pierce™ Silver Stain Kit (Thermo Fisher Scientific, catalog number 24612).

### Statistics and reproducibility

Sample sizes and statistical tests are noted in the figure captions. Illumina sequencing was performed with three independent replicates for the LASSO, NEBNext, and *riboPOOLs* samples. One *riboPOOLs* library failed quality control due to insufficient final RNA concentration, reducing the number of *riboPOOLs* replicates to two. All other assays were performed with three independent replicates for each sample. No data were excluded from the analysis. All experiments were performed multiple times and showed consistent results, as reported in the manuscript.

## Acknowledgements

E.K. acknowledges funding by the Federal Ministry of Research, Technology and Space (BMFTR) in the program NanoMatFutur (grant no. 13XP5098). We thank Dr. Andreas Dahl, Dr. Maximilian Krause, Dr. Mathias Lesche, and Dr. Julieta Aprea from the Dresden Concept Genome Center for helpful discussions regarding RNA-seq data analysis. We also thank Dr. Syuan-Ku Hsiao for his assistance with polymer synthesis. We thank Dr. Hartmut Komber for assistance with NMR measurements. We thank Dr. Manfred Maitz for valuable discussions regarding thrombin pulldown and quantification.

## Author contributions

E.K. conceived the project. S.K.S. carried out polymer synthesis, pulldown experiments, system optimizations, and data analysis with input from E.K. and K.G. K.G. performed initial studies on polymer pulldown through DNA crosslinking. Y.-H.P. performed the confocal microscopy imaging. S.K.S. and E.K. wrote the manuscript with input from all coauthors.

## Data availability

All data supporting the findings of this study are provided in the article and its Supplementary Information.

