## Supplementary Information for "LASSO: versatile and selective biomolecule pulldown with combinatorial DNA-crosslinked polymers"

#### 1 Supplementary Procedures

***Supplementary Procedure 1: General LASSO capture and release***

1. **Reagents**

- 0.5% (w/v) DNA-functionalized polymer with 100 µM (**$P_{10}$**) or 200 µM (**$P_{20}$**) maximum anchor strand DNA (Supplementary Data 1, strand ID 1) in 1x TE buffer
- 10x Hybridization buffer (**HB**): 100 mM Tris, 10 mM EDTA, 1.5 M NaCl, pH 8.0
- **$H_2O$**
- Catcher strand library DNA (**CSL**, mixed in equimolar ratio) in 1x TE buffer
- 100 µM combinatorial crosslinker library with 64 unique overlap sequences (**CCL-64**, mixed in equimolar ratio) in 1x TE buffer (Supplementary Data 1, strand IDs 9 and 10)
- Release strand library DNA (**RSL**, mixed in equimolar ratio) in 1x TE buffer
- Biomolecule **target** in buffer of choice (e.g. $H_2O$, TE buffer, PBS buffer)

2. **Target capture**

A) To capture desired target (ssDNA, RNA, or protein) select the appropriate CSL and target and mix the following components together:

| Component | Final Concentration |
|---|---|
| $P_{10}$ or $P_{20}$ | 0.05% w/v (max. 10 µM or 20 µM anchor stands) |
| CSL | Up to 0.1 molar equivalents of anchor strands |
| CCL-64 | 0.8 molar equivalents of anchor strands |
| Target* | 0.3 molar equivalents of CSL |
| HB | To 1x concentration |
| $H_2O$ | Up to final desired volume |

*Target can be added before annealing if capturing nucleic acids, or after annealing if capturing temperature-sensitive molecules (e.g. proteins)

B) Vortex thoroughly, then anneal as follows:
(1) Heat to 95 °C for 3 min,
(2) Instant cool from 95 °C to 80 °C,

(3) Hold at 80 °C for 2 min,
(4) Cool from 80 °C to 65 °C at −1.5 °C $min^{-1}$,
(5) Cool from 65 °C to 37 °C at −2.8 °C $min^{-1}$,
(6) Hold at preferred temperature (e.g. 20 °C or 4 °C) until use.

C) If capturing target after annealing, add target and bind for 30 minutes with light shaking at preferred temperature.

D) Centrifuge at 17,000 xg for 30 minutes at preferred temperature.

E) The supernatant is then retrieved if target depletion is desired; optionally, the target can be released from the polymer pellet.

**3. Target release**

A) Remove supernatant, leaving polymer pellet intact at bottom of tube.

B) Optionally, wash polymer pellet 3x with 1x TE buffer + 150 mM NaCl by gently adding the buffer without disturbing the pellet, incubating for 30 seconds, then removing the same volume of buffer.

C) Add 1.5x molar excess of RSL over the CSL concentration and resuspend the pellet with vigorous pipetting.

D) Incubate for at least two hours with light shaking at preferred temperature.

E) Centrifuge at 17,000 xg for 30 minutes at preferred temperature.

F) The supernatant containing the released target is retrieved.

***Supplementary Procedure 2: LASSO ssDNA/RNA capture***

**1. Reagents**

- 0.5% (w/v) DNA-functionalized polymer with 100 µM (**$P_{10}$**) maximum anchor strand DNA (Supplementary Data 1, strand ID 1)
- 10x Hybridization buffer (**HB**): 100 mM Tris, 10 mM EDTA, 1.5 M NaCl, pH 8.0
- **$H_2O$**
- 5 µM catcher strand library DNA (**CSL**, mixed in equimolar ratio) in 1x TE buffer (Supplementary Data 1, strand ID 14 for DNA, strand IDs 18 and 19 for RNA)
- 100 µM combinatorial crosslinker library with 64 unique overlap sequences (**CCL-64**, mixed in equimolar ratio) in 1x TE buffer (Supplementary Data 1, strand IDs 9 and 10)
- 0.5 µM **target** single-stranded DNA or RNA in 1x TE buffer (Supplementary Data 1, strand ID 13 for DNA, strand ID 17 for RNA)

#### 2. Nucleic acid capture

A) To capture specific target nucleic acids (ssDNA or RNA), select the appropriate CSL and target and mix the following components together:

| Component | Volume (µL) | Final Concentration |
|---|---|---|
| $P_{10}$ | 5 | 0.05% w/v (10 µM anchor strands) |
| CSL | 1.5 | 150 nM |
| CCL-64 | 4 | 8 µM |
| Target | 5 | 50 nM |
| HB | 5 | 1x |
| $H_2O$ | 29.5 | |
| Total | 50 | |

B) Vortex thoroughly, then anneal as follows:
(1) Heat to 95 °C for 3 min,
(2) Instant cool from 95 °C to 80 °C,
(3) Hold at 80 °C for 2 min,
(4) Cool from 80 °C to 65 °C at −1.5 °C $min^{-1}$,
(5) Cool from 65 °C to 37 °C at −2.8 °C $min^{-1}$,
(6) Hold at 20 °C until use.

C) Centrifuge samples at 17,000 xg for 30 minutes at room temperature.

***Supplementary Procedure 3: LASSO rRNA depletion***

#### 1. Reagents

- 0.5% (w/v) DNA-functionalized polymer with 200 µM (**$P_{20}$**) maximum anchor strand DNA (Supplementary Data 1, strand ID 1)
- 10x Hybridization buffer (**HB**): 100 mM Tris, 10 mM EDTA, 1.5 M NaCl, pH 8.0
- Nuclease-free **$H_2O$**
- 0.5 M **EDTA** pH 8.0
- ~100 µM catcher strand library DNA (**CSL**) in nuclease-free $H_2O$ (Supplementary Data 1, mix of 0.45 µM each of strand IDs 22 – 210 and 2.25 µM each of strand IDs 211 – 216)
- 100 µM combinatorial crosslinker library with 64 unique overlap sequences (**CCL-64**, mixed in equimolar ratio) in 1x TE buffer (Supplementary Data 1, strand IDs 9 and 10)
- 1 µg/µL human HeLa cell **total RNA** in $H_2O$
- DNase I (NEB, catalog number M0303S)
- 10x DNase I buffer

- Monarch® Spin RNA Cleanup Kit (NEB, catalog number T2050S)

#### 2. rRNA capture

A) To deplete rRNA, mix the following components together on ice:

| Component | Volume (µL) | Final Concentration |
|---|---|---|
| $P_{20}$ | 25 | 0.05% w/v (20 µM anchor strands) |
| CSL | 3.3 | 1.32 µM, 5000 ng |
| CCL-64 | 40 | 16 µM |
| Total RNA | 1 | ~100 nM, 1000 ng |
| HB | 25 | 1x |
| $H_2O$ | 155.7 | |
| Total | 250 | |

B) Vortex thoroughly, then anneal as follows in a pre-heated thermocycler:
(1) Heat to 95 °C for 3 min,
(2) Instant cool from 95 °C to 80 °C,
(3) Hold at 80 °C for 2 min,
(4) Cool from 80 °C to 65 °C at −1.5 °C $min^{-1}$,
(5) Cool from 65 °C to 37 °C at −2.8 °C $min^{-1}$,
(6) Hold at 4 °C until use.

C) Centrifuge samples at 17,000 xg for 30 minutes at 4 °C.

D) Transfer 230 µL of supernatant to a clean tube on ice.

#### 3. RNA purification

A) Purify the rRNA-depleted RNA library using the Monarch® Spin RNA Cleanup Kit, eluting into a final volume of 30 µL $H_2O$.

B) To degrade any remaining oligonucleotides, mix the following components together on ice:

| Component | Volume (µL) |
|---|---|
| rRNA-depleted RNA library | 30 |
| DNase I | 1 |
| 10x DNase I buffer | 3.5 |
| $H_2O$ | 0.5 |
| Total | 35 |

C) Incubate at 37 °C for 10 minutes.

D) Add EDTA to a final concentration of 5 mM and purify the final RNA library using the Monarch® Spin RNA Cleanup Kit.

##### *Supplementary Procedure 4: LASSO thrombin capture and release*

**1. Reagents**

- 0.5% (w/v) DNA-functionalized polymer with 100 µM (**$P_{10}$**) maximum anchor strand DNA (Supplementary Data 1, strand ID 1)
- 10x Hybridization buffer (**HB**): 100 mM Tris, 10 mM EDTA, 1.5 M NaCl, pH 8.0
- **1% BSA buffer**: 10 mM Tris, 1 mM EDTA, 150 mM NaCl, 1% (w/v) bovine serum albumin (BSA), pH 8.0
- **0.1% BSA buffer**: 10 mM Tris, 1 mM EDTA, 150 mM NaCl, 0.1% (w/v) BSA, pH 8.0
- **$H_2O$**
- 10 µM catcher strand library DNA (**CSL**, mixed in equimolar ratio) in 1x TE buffer (Supplementary Data 1, strand IDs 229 and 231)
- 100 µM combinatorial crosslinker library with 64 unique overlap sequences (**CCL-64**, mixed in equimolar ratio) in 1x TE buffer (Supplementary Data 1, strand IDs 9 and 10)
- 10 µM release strand library DNA (**RSL**, mixed in equimolar ratio) in 1x TE buffer (Supplementary Data 1, strand IDs 230 and 232)
- 0.6 µM human thrombin protein **target** in 1% BSA buffer

**2. Thrombin capture**

A) To capture thrombin, mix the following components together:

| Component | Volume (µL) | Final Concentration |
|---|---|---|
| $P_{10}$ | 5 | 0.05% w/v (10 µM anchor strands) |
| CSL | 2.5 | 500 nM |
| CCL-64 | 4 | 8 µM |
| HB | 4.5 | 1x |
| $H_2O$ | 29 | |
| Total | 45 | |

B) Vortex thoroughly, then anneal as follows:
(1) Heat to 95 °C for 3 min,
(2) Instant cool from 95 °C to 80 °C,
(3) Hold at 80 °C for 2 min,
(4) Cool from 80 °C to 65 °C at $-1.5$ °C $min^{-1}$,

(5) Cool from 65 °C to 37 °C at −2.8 °C $min^{-1}$,
(6) Hold at 20 °C until use.

C) Add 5 µL of 0.6 µM thrombin (in 1% BSA buffer) to bring final volume to 50 µL.

D) Incubate sample for 30 minutes with light shaking at room temperature.

E) Centrifuge at 17,000 xg for 30 minutes at room temperature.

**3. Thrombin release**

A) Remove supernatant, leaving polymer pellet intact at bottom of tube.

B) Wash polymer pellet 3x with 200 µL of 0.1% BSA buffer by gently adding the buffer without disturbing the pellet, incubating for 30 seconds, then removing the same volume of buffer.

C) Add 3.75 µL of RSL and resuspend the pellet with vigorous pipetting.

D) Incubate for two hours with light shaking at room temperature.

E) Centrifuge at 17,000 xg for 30 minutes at room temperature.

F) The supernatant containing the released thrombin is retrieved for analysis.

### 2 Supplementary Methods

#### 2.1 Polyacrylamide gel electrophoresis (PAGE)

The binding capacity of $\mathbf{P_{10}}$ and $\mathbf{P_{20}}$ and the stability of $\mathbf{P_{10}}$ over long-term storage were determined by PAGE. Samples were prepared in 1x TE buffer (10 mM Tris, 1 mM EDTA, pH 8.0) with a final concentration of 150 mM NaCl. To test the binding capacity, either 0.0025% (w/v) of $\mathbf{P_{10}}$ (maximum 0.5 µM anchor strands) or 0.00125% (w/v) of $\mathbf{P_{20}}$ (maximum 0.5 µM anchor strands) was used. To test the stability of $\mathbf{P_{10}}$, 0.0033% (w/v) of a $\mathbf{P_{10}}$ stock (maximum 0.5 µM anchor strands) that had been stored at −20 °C for seven years was used. The original concentration of anchor strands after synthesis was quantified as 75 µM for the 0.5% (w/v) $\mathbf{P_{10}}$ stock. The binding efficiency test target strand (Supplementary Data 1, strand ID 2) was added at a series equivalent from 0.2x–2.0x the amount of anchor strands for all samples. The samples were annealed according to the LASSO protocol (Supplementary Procedure 1). Native polyacrylamide gels were prepared using a 40% (w/w) acrylamide/bis-acrylamide (19:1) stock. The annealed samples were loaded into a 15% native polyacrylamide gel and run in 0.5x TBE buffer (50 mM Tris, 45 mM boric acid, 0.5 mM EDTA, pH 8.3) at 100V on a XCell SureLock Mini-Cell Electrophoresis System using a Consort™ EV265 Electrophoresis power supply (Thermo Fisher Scientific). Gels were stained with SYBR™ Gold. Gel scans (using a blue laser excitation at 473 nm) were recorded on a Typhoon FLA 9500 scanner (GE Healthcare Life Sciences) at a 50 µm/pixel resolution with the accompanying software (v1.0).

#### 2.2 Nuclear magnetic resonance spectroscopy

Nuclear magnetic resonance (NMR) spectroscopy was performed on samples of unpurified polymers prepared at a 1:8 dilution in $D_2O$. $^1H$ NMR spectra were recorded at 30–32 °C on a 500 MHz spectrometer (Bruker) with a 2 second acquisition time and 32 transients. Chemical shifts (δ) are reported in parts per million (ppm) downfield from tetramethyl silane (TMS). The $^1H$ NMR shifts are relative to the residual hydrogen peak of $D_2O$ (4.79 ppm).

#### 2.3 Optimization of the LASSO protocol

Fluorescence images of all Eppendorf tubes (in the Cy5 channel; excitation at 635 nm) were recorded before and after centrifugation on a Typhoon FLA 9500 scanner (GE Healthcare Life Sciences) at a 50 µm pixel size with the accompanying software (v1.0). Image analysis of fluorescence was performed in Fiji (v1.54m). Capture efficiency was determined by comparing the fluorescence intensity in the supernatant to a DNA-only control sample after centrifugation.

##### 2.3.1 Confocal microscopy

Samples were prepared with 0.05% (w/v) $\mathbf{P_{10}}$ and 80% of the anchor strand concentration of each combinatorial crosslinker library (CCL) containing 0 (uncrosslinked control), 1, 4, 16, 64 or 256 unique crosslinkers (Supplementary Data 1, strand IDs 3 – 12) in 1x TE buffer and 150 mM NaCl. The samples were annealed according to the LASSO protocol (Supplementary Procedure 1). DNA in the sample was stained with SYBR™ Gold for visualization. Confocal images were acquired at 100x magnification on an Andor Dragonfly confocal microscope (Oxford Instruments).

##### 2.3.2 Crosslinker diversity

The number of unique crosslinkers in each combinatorial crosslinker library (CCL) was tested for its effect on the phase separation of the polymer and capture efficiency on LASSO. Samples were prepared with 0.02% (w/v) $\mathbf{P_{10}}$, 75% of the anchor strand concentration of each combinatorial crosslinker library (CCL) containing 0 (uncrosslinked control), 1, 4, 16, 64 or 256 unique crosslinkers (Supplementary Data 1, strand IDs 3 – 12), 100 nM catcher strand (Supplementary Data 1, strand ID 14), and 50 nM fluorescent ssDNA oligonucleotide target (Supplementary Data 1, strand ID 13) in 1x TE buffer and 150 mM NaCl. The samples were annealed and centrifuged according to the LASSO protocol (Supplementary Procedure 1).

##### 2.3.3 Polymer concentration

The polymer concentration was tested for its effect on the phase separation of the polymer and capture efficiency on LASSO. Samples were prepared with 0.01–1% (w/v) $\mathbf{P_{10}}$, 90% of the anchor strand concentration of CCL-64 (Supplementary Data 1, strand IDs 9 and 10), 150 nM catcher strand (Supplementary Data 1, strand ID 14), and 50 nM fluorescent ssDNA oligonucleotide target (Supplementary Data 1, strand ID 13) in 1x TE buffer and 150 mM NaCl. The samples were annealed and centrifuged according to the LASSO protocol (Supplementary Procedure 1).

##### 2.3.4 Crosslinker concentration

The concentration of crosslinkers was tested for its effect on the phase separation of the polymer and capture efficiency on LASSO. Samples were prepared with 0.05% (w/v) $\mathbf{P_{10}}$, 10%–100% of the anchor strand concentration of CCL-64 (Supplementary Data 1, strand IDs 9 and 10), 150 nM catcher strand (Supplementary Data 1, strand ID 14), and 50 nM fluorescent ssDNA oligonucleotide target (Supplementary Data 1, strand ID 13) in 1x TE buffer and 150 mM NaCl. The samples were annealed and centrifuged according to the LASSO protocol (Supplementary Procedure 1).

#### 2.4 Testing adsorption of RNA to DNA-grafted poly(acrylamide-coacrylic acid) during methanol precipitation

Cytoplasmic RNA was mixed with DNA-functionalized polymers (RNA + Polymer) or with an equivalent volume of 1x TE buffer (RNA only) with varying pH and NaCl concentrations and subjected to methanol precipitation as described in ref. [1]. Absorbance readings at 230, 260, and 280 nm were taken in a spectrophotometer to determine RNA concentration in the supernatant.

### 3 Supplementary Notes

***Supplementary Note 1: Effect of crosslinker diversity on polymer agglomeration behavior***

As shown in Figure 1c in the main text, the agglomeration and sedimentation behavior of the polymer strongly depends on the diversity of the CCL. The striking difference between CCL-64-crosslinked polymer versus higher or lower complexity libraries demands an explanation.

As reported previously,[2] the polymer used in this experiment, $\mathbf{P_{10}}$, has a molecular weight of approximately 2.3 MDa, and each chain is functionalized with 28 anchor strands, on average. The statistical simulation from the same study reveals that 28 anchor strands would require at least 64 distinct splint pairs to suppress intra-molecular bonds effectively (Fig. 2a from ref. 2). This requirement explains why large polymer agglomerates are not observed when using CCL-1, CCL-4, and CCL-16, as opposed to CCL-64, which produces large agglomerates.

The simulation, however, does not explain why CCL-256 does not yield the same or even superior agglomeration and sedimentation behavior than CCL-64. We surmise that this effect is due to the expected reduction in crosslinking kinetics: As CCL-256 has a 4x higher number of distinct crosslinker pairs, each of the individual crosslinkers is present at a 4x lower concentration. DNA hybridization follows second-order reaction kinetics. Therefore, the 4x lower concentration must result in 16x slower crosslinking kinetics, and thus CCL-256 is expected to require 16x longer for polymer chains to agglomerate and reach the same size as CCL-64-crosslinked polymers.

In addition to the primary "dilution effect" on crosslinking kinetics, one would also expect slightly less favorable binding efficiencies in CCL-256 versus CCL-64 at thermodynamic equilibrium. To quantify this effect, we performed nearest-neighbor thermodynamic calculations[3] using a representative 14-nt overlap sequence (`TTAGTTAGTATCCC`) and its complement (`GGGATACTAACTAA`) with experimental parameters of 150 mM monovalent cations, 25°C, 8 µM total anchor strand concentration, and 0.8 equivalents of crosslinkers. The calculations reveal that approximately 6% of CCL-64 overlap domains stay unpaired at equilibrium (each overlap domain having 100 nM concentration). This value increases to 12% for CCL-256 (where each overlap domain has 25 nM concentration). CCL-256 is therefore predicted to yield a slightly lower number of bound crosslinkers at the same total crosslinker concentration.

***Supplementary Note 2: Optimization of polymer concentration***

The concentration of the polymer is vital for the formation of individual polymer agglomerates that are dense enough to properly pellet and efficiently capture the target. If the concentration is higher than the critical gelation concentration of 0.2% (w/v),[2] the crosslinked polymers will form an extended polymer gel that cannot be easily compressed into a compact pellet by centrifugation. However, if the concentration is too low, the polymer chains do not efficiently cross-link and target capture is reduced. We found that a polymer concentration of 0.05% (w/v) exhibits the highest capture efficiency of a fluorescent DNA oligonucleotide target while forming a sufficiently compact pellet (Supplementary Fig. 4).

### 4 Supplementary Figures

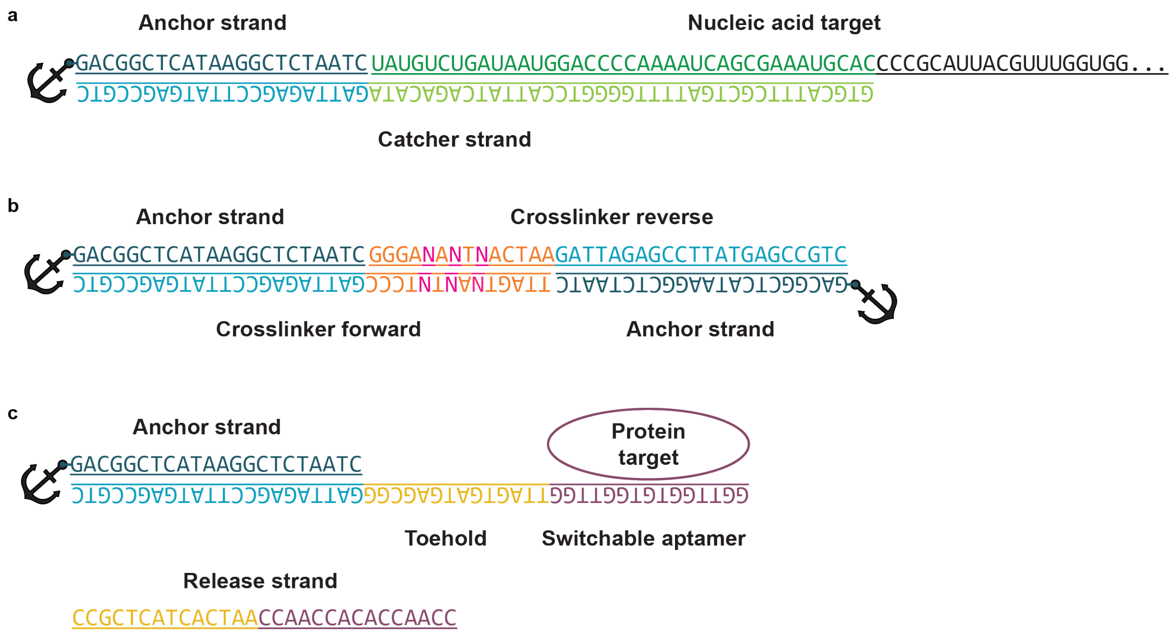


***Supplementary Figure 1. Schematic representation of possible DNA complexes on LASSO. a)*** *Capture of nucleic acids on a catcher strand containing a target-specific binding region and a region complementary to the polymer-bound anchor strand.* ***b)*** *Crosslinking between polymer chains with a combinatorial crosslinker library (CCL). Mixed N bases in the overlap are colored in magenta.* ***c)*** *Capture of a protein target on a switchable aptamer catcher strand. The target can be released via toehold-mediated strand displacement (TMSD) through the addition of a release strand. Anchor strands are conjugated to the polymer (depicted as anchor symbols).*

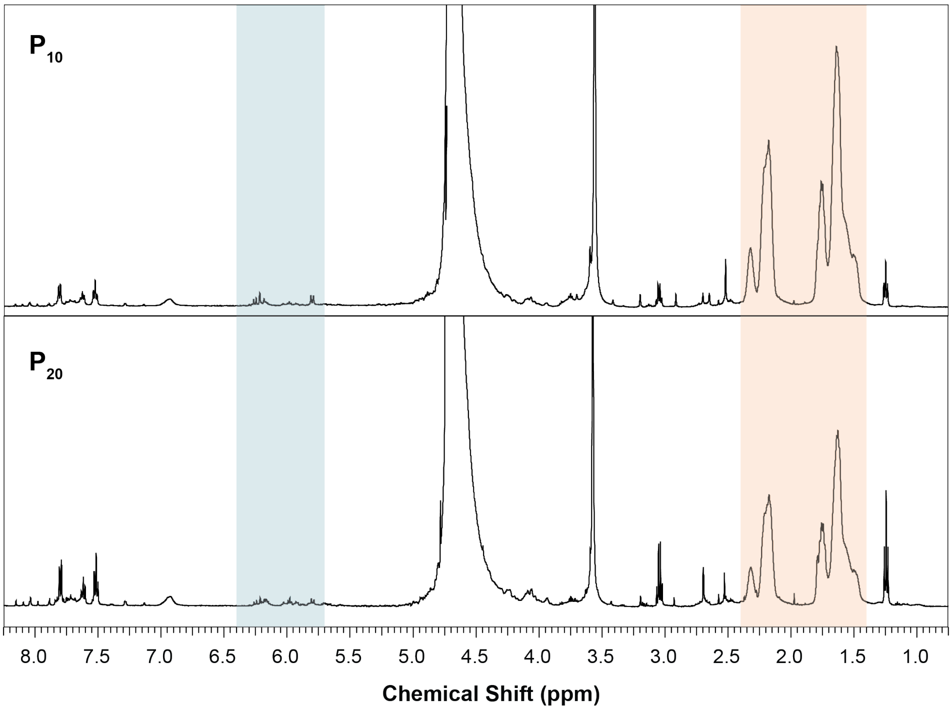


***Supplementary Figure 2. $^1$H-NMR spectra of $P_{10}$ and $P_{20}$ in $D_2O$ prior to methanol purification.*** The conversion percentage is determined by measuring the ratio of free residual acrylamide monomer protons (δ ~5.7–6.4; blue) to polymer backbone protons (δ ~1.4–2.4; orange). $\mathbf{P_{10}}$ showed 99% conversion; $\mathbf{P_{20}}$ showed 98% conversion.

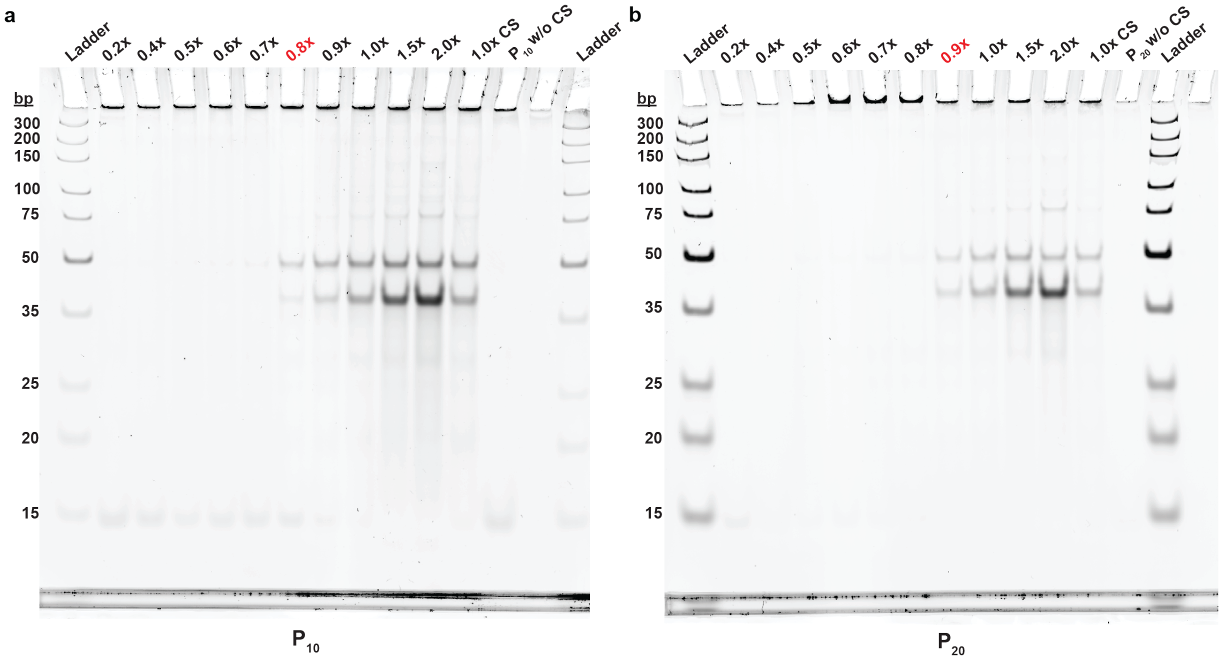


***Supplementary Figure 3. DNA binding capacity test on $P_{10}$ and $P_{20}$.*** *The polymer was mixed with catcher strands (CS) complementary to the anchor strand at a concentration range from 0.2x to 2.0x, where 1x is defined as 100% of the maximum binding capacity.* ***a) $P_{10}$*** *saturated with CS at 0.8x, indicating the available concentration of anchor strands was approximately 80 µM in a 0.5% (w/v) solution.* ***b) $P_{20}$*** *saturated with CS at 0.9x, indicating the available concentration of anchor strands was approximately 170 µM in a 0.5% (w/v) solution.*

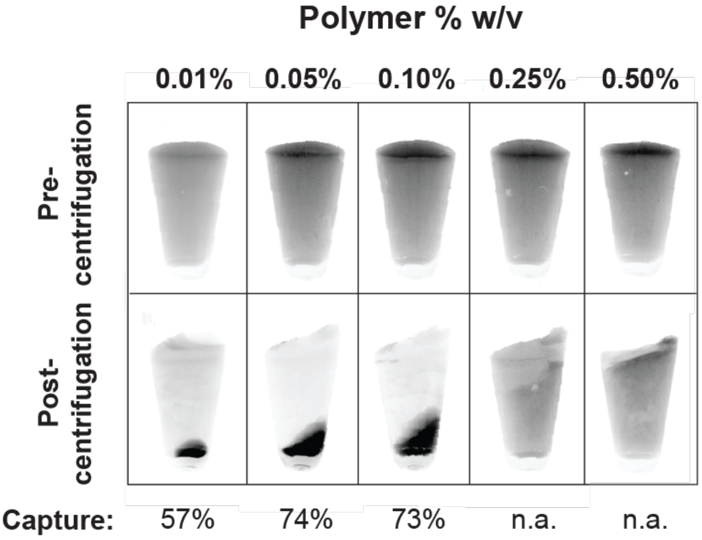


***Supplementary Figure 4. Optimization of polymer concentration.*** *Capture of a fluorescent single-stranded DNA oligonucleotide on* $\boldsymbol{P_{10}}$ *at different concentrations reveals the highest capture efficiency at 0.05% (w/v). Concentrations higher than the critical gelation concentration of 0.2% (w/v) start to form an extended polymer gel that cannot be pelleted.*

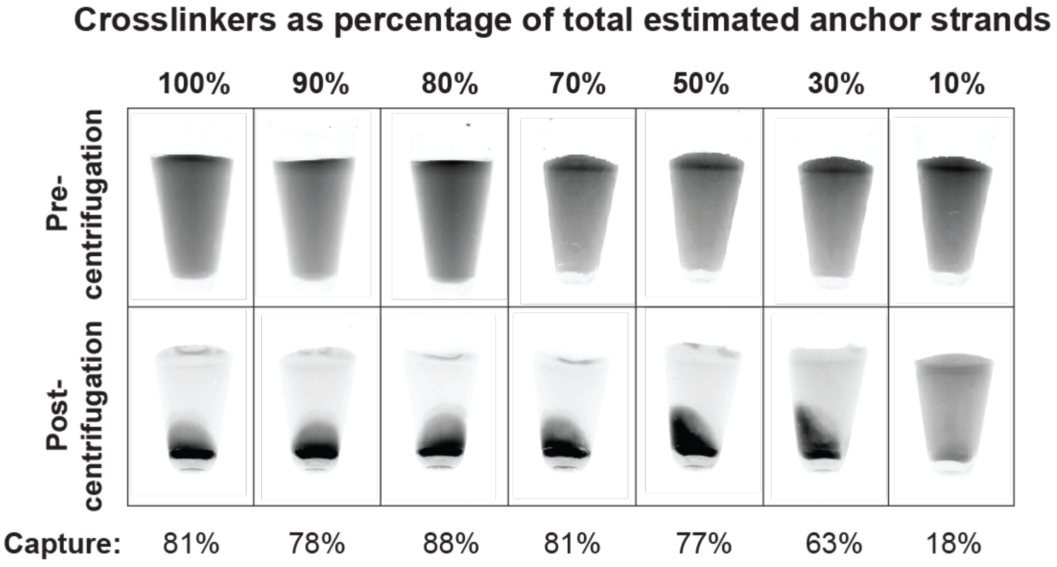


***Supplementary Figure 5. Optimization of crosslinker concentration.*** *Capture of a fluorescent single-stranded DNA oligonucleotide on 0.05% (w/v)* $\boldsymbol{P_{10}}$ *at different CCL-64 concentrations reveals the highest efficiency of capture at 80% of the total estimated anchor strand concentration.*

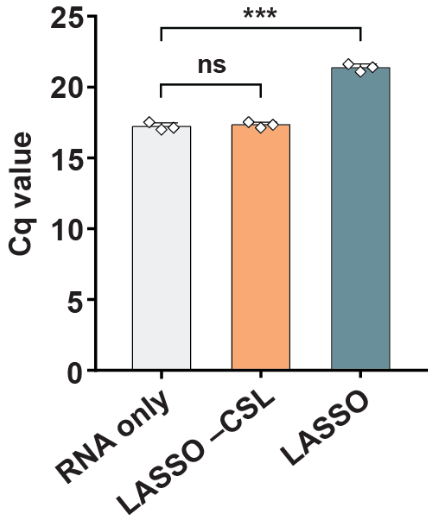


***Supplementary Figure 6. Cq values for RT-qPCR of SARS-CoV-2 N-gene RNA after depletion with LASSO.*** *Data are shown as mean ± s.d. (n = 3 independent experiments). Statistical analysis was performed using an unpaired two-tailed t-test; ns, non-significant; ***p < 0.001.*

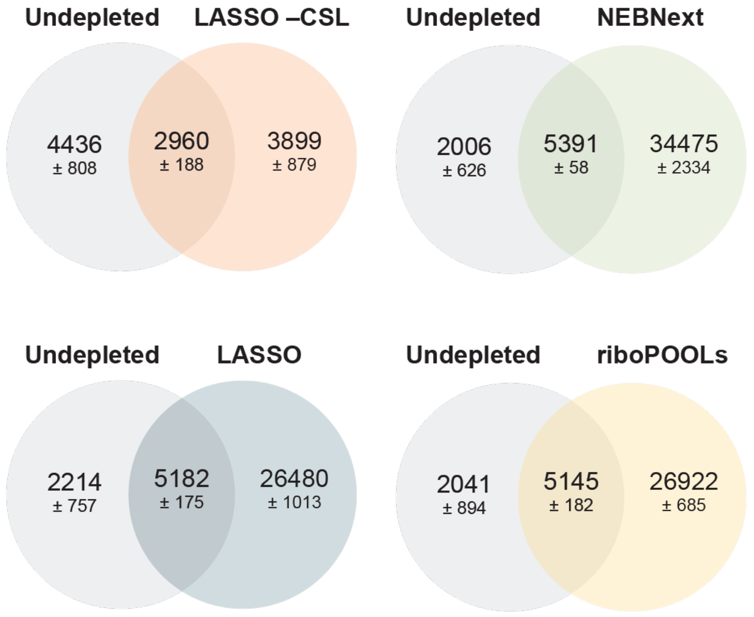


***Supplementary Figure 7. Comparison of RNA library composition after rRNA depletion.*** *Venn diagrams for all transcripts with TPM > 1 in rRNA-depleted samples (and the –CSL control) versus the undepleted total RNA sample.*

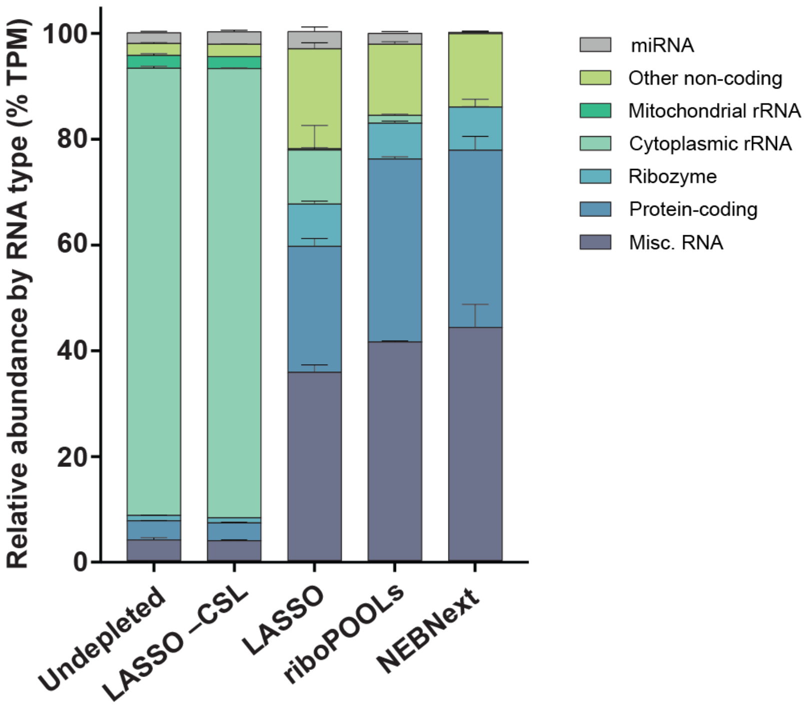


***Supplementary Figure 8. Biotype distribution including rRNA reads.*** *All RNA reads were mapped to their respective Ensembl biotype annotation, normalized to transcript per million (TPM), and grouped by type.*

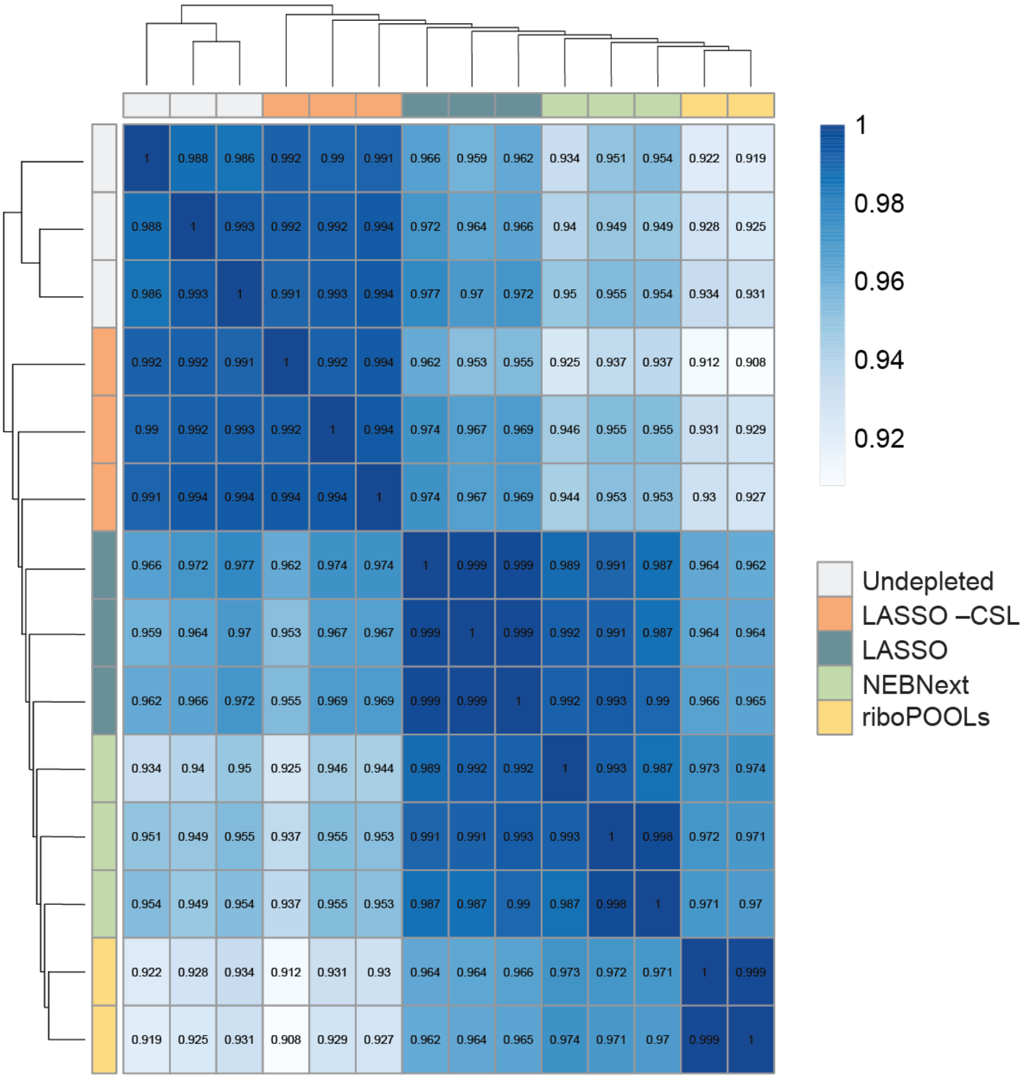


***Supplementary Figure 9. Comparison of Pearson correlation coefficients between methods for rRNA depletion (excluding rRNA depletion targets).***

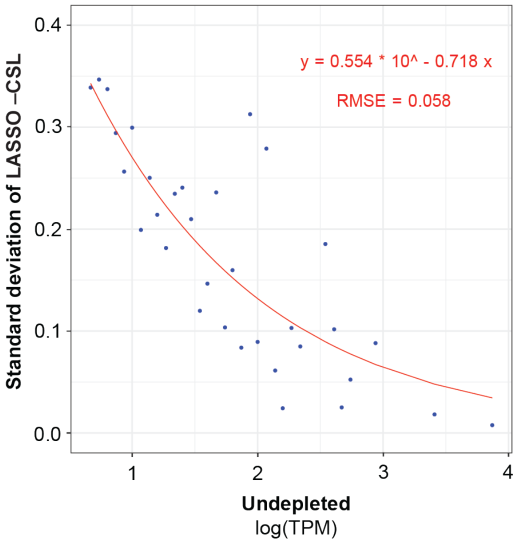


***Supplementary Figure 10. Modeling of expected standard deviations for rRNA-depleted samples.*** *Outliers in Figure 3d were determined using the exponential model derived in this plot. See Methods section "RNA sequencing (RNA-seq)" for details.*

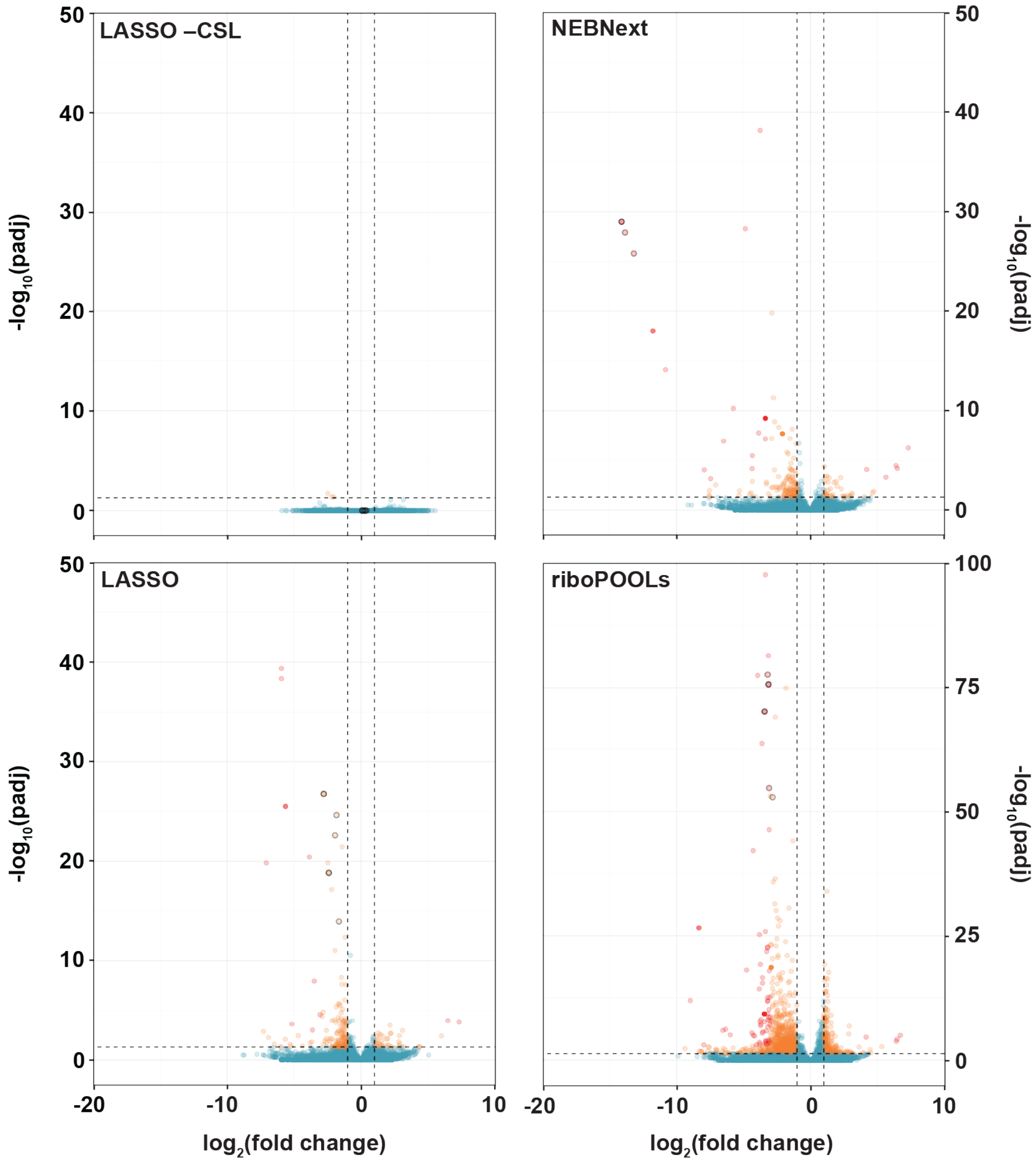


***Supplementary Figure 11. Differential transcript detection from RNA-seq.*** *Volcano plots showing $log_2$ fold changes in detected transcript abundance between rRNA depletion methods, excluding rRNA transcripts. Transcripts with padj < 0.05 and absolute $log_2$ fold change > 1 (orange) or padj < 0.001 and absolute $log_2$ fold change > 3 (red) are considered significantly affected or extremely affected, respectively. Datapoints with black outlines indicate transcripts with high sequence similarity (>90%) to rRNA and are identical to the outlined datapoints in Figure 3d.*

**a**

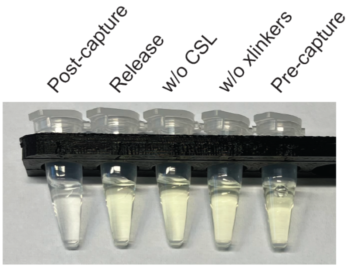


**b**

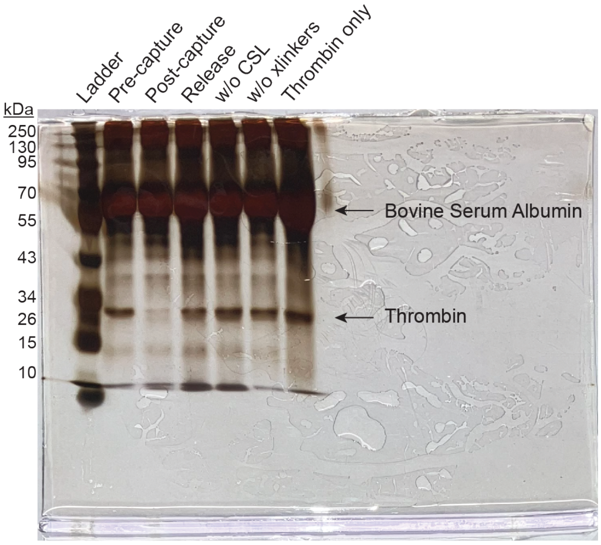


***Supplementary Figure 12*. Raw images for thrombin capture and release on LASSO. a)** *Raw image of tubes containing supernatant samples after cleavage of S-2238 substrate by thrombin. Samples were incubated for 20 minutes at 37°C.* ***b)*** *Raw image of supernatant samples on silver-stained 12.5% SDS-PAGE gel. The buffer contains BSA (66 kDa), a common additive for protein stabilization. For details, see Supplementary Procedure 4.*

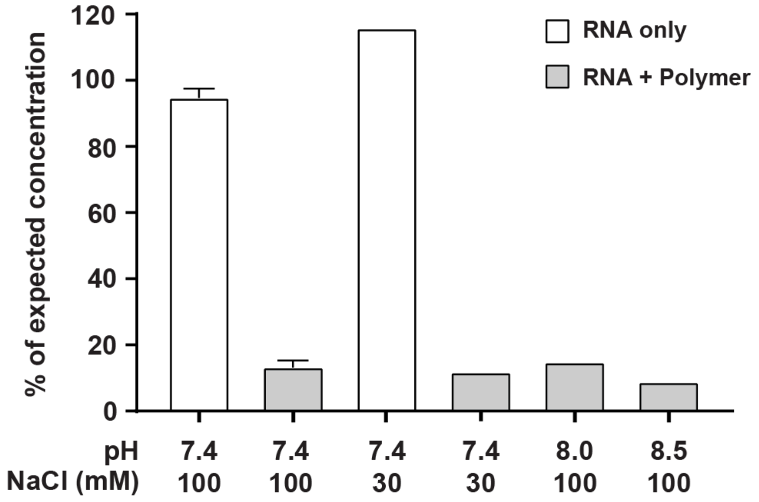


***Supplementary Figure 13. Nonspecific RNA adsorption on DNA-grafted poly(acrylamide-coacrylic acid) during methanol precipitation.*** *n = 2 for samples at pH 7.4, 100 mM NaCl; n = 1 for all other samples.*

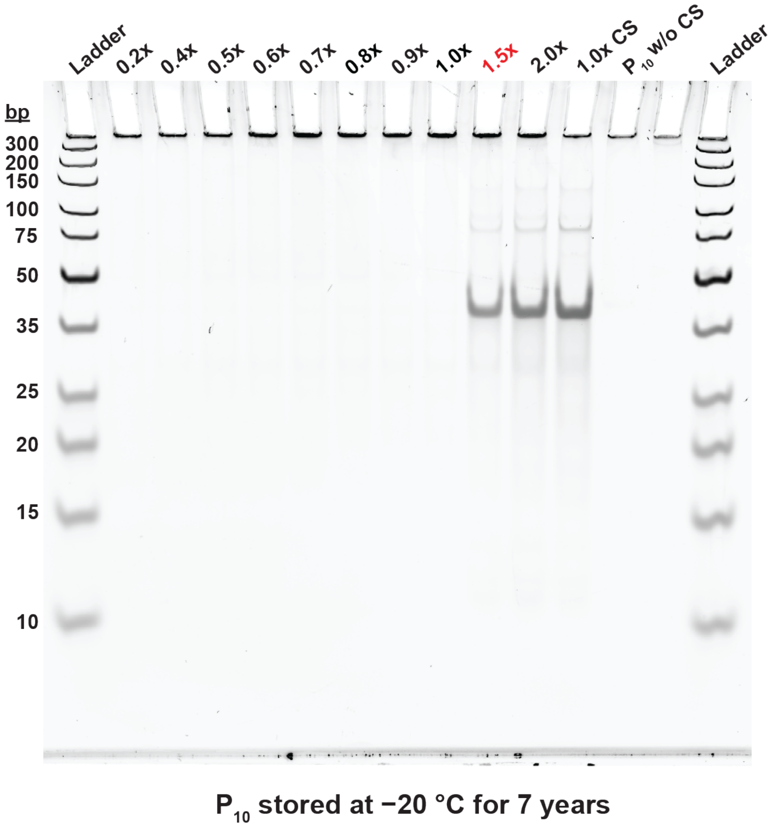


***Supplementary Figure 14. Long-term stability test of $P_{10}$.*** *The polymer had been stored at −20 °C for seven years. The original concentration of anchor strands after synthesis was quantified as 75 µM for a 0.5% (w/v) stock solution. The polymer was mixed with catcher strands (CS) complementary to the anchor strand at a concentration range from 0.2x to 2.0x, where 1x is defined as 100% of the original binding capacity.* ***$P_{10}$*** *saturated with catcher strands (CS) at 1.5x, indicating the available concentration of anchor strands was approximately 75 µM in a 0.5% (w/v) stock solution after long-term storage, or 100% of the original binding capacity.*

#### 5 Supplementary Tables

***Supplementary Table 1. Cost calculation of LASSO.*** *The cost is calculated per pmol target for general capture and release (for example, a single nucleic acid sequence or protein) and for rRNA depletion using a full CSL on 1 µg of total RNA. The purchase price per reaction for the two commercial rRNA depletion kits is also given.*

| Materials/Reagents | Unit | Unit cost | Cost for general capture and release | Cost for rRNA depletion |
|---|---|---|---|---|
| Anchor strand | 1 nmol | $ 0.07 | $ 0.02 | $ 0.35 |
| DNA crosslinkers | 1 nmol | $ 0.09 | $ 0.02 | $ 0.36 |
| DNA catcher strands | 1 nmol | $ 0.15/$ 0.36 | $ 0.00 | $ 0.11 |
| DNA release strands | 1 nmol | $ 0.10 | $ 0.00 | - |
| Hybridization buffer | 1 mL | $ 0.06 | $ 0.00 | $ 0.00 |
| DNase I | 1 unit | $ 0.07 | - | $ 0.14 |
| **Total cost for LASSO pulldown assay** | | | **~ $ 0.04** | **~ $ 0.96** |
| **Cost for NEBNext rRNA depletion kit v2** | | | **-** | **~ $ 51.00** |
| **Cost for riboPOOLs rRNA depletion kit** | | | **-** | **~ $ 46.00** |

***Supplementary Table 2. Comparison between LASSO and commercial kits for rRNA depletion.***

| Protocol | Protocol Time* | Pros | Cons |
|---|---|---|---|
| LASSO | 1 hour | • Minimal bias and off-target capture<br>• Easy to use (mix and spin)<br>• Inexpensive and highly scalable<br>• Long shelf life | • Moderate depletion efficiency (~86%) |
| riboPOOLs | 1 hour | • High depletion efficiency (~97%)<br>• Easy to use (mix and pull)<br>• Automation-friendly | • Introduces substantial bias<br>• Expensive |
| NEBNext | 1.5 hours | • Exceptional depletion efficiency (>99%) | • Introduces some bias<br>• Short shelf life<br>• Expensive<br>• Multi-step protocol |

*Net time of the depletion step, not including any additional purification (for example, with SPRI beads or a silica spin column).
